# Phenotypic lags influence rapid evolution throughout a drought cycle

**DOI:** 10.1101/2023.08.18.553905

**Authors:** Haley A. Branch, Daniel N. Anstett, Amy L. Angert

## Abstract

Climate anomalies pose strong selection which can lead to rapid evolution. These global mean trends occur on a backdrop of interannual variability that might weaken or even reverse selection. However, the impact of climatic interannual variability on rapid evolution is rarely considered. We study evolution through a seven-year period encompassing a severe drought across 12 populations of *Mimulus cardinalis* (scarlet monkeyflower). Plants were grown in a common greenhouse environment under wet and dry treatments, where specific leaf area and date of flowering were measured. We compare the ability of different climate metrics to explain the rapid evolution of trait values, examining different time-periods, including the collection year, prior years, and cumulative metrics across sequential years. We find that anomalies in mean annual precipitation best describe rapid evolution over our study period. Past climates, of one-to two-years ago, are often related to trait values in a conflicting direction to collection-year climate. Uncovering these complex climatic impacts on evolution is critical to better predict and interpret the impacts of climate change.

## Introduction

Climate change may require species to quickly evolve adaptations as environments become unsuitable (Franks et al. 2007; Catullo et al. 2019). It is now well established that evolution frequently occurs on contemporary timescales (Reznick et al. 1990; Agrawal et al. 2012; Grant and Grant 2014). However, it remains unclear whether rapid evolution during climate change will be sufficient to reverse demographic declines, a process known as evolutionary rescue (Gomulkiewicz and Holt 1995; Bell and Gonzalez 2011; Bell 2017). Rate of adaptation is a function of extrinsic and intrinsic factors, where extrinsic factors refer to the strength of selection imposed by novel environments and intrinsic factors refer to population capacity to respond to selection. While studies seeking to understand rapid evolution often consider the impact of climate (extrinsic), they rarely consider the legacy of selection during prior years and how it may impact a species’ intrinsic ability to adapt. These legacies can be particularly important for long-lived species where the impact of selection might not be fully realized until years after the selection event.

Strength of selection is determined at least in part by the intensity and length of exposure to anomalous environments (i.e., deviations from historical baselines). These deviations can also vary from year to year, resulting in perennial populations where phenotypes observed during a single year are the combined evolutionary product across multiple years. Across years, selection can be a consistent press perturbation or it can relax and even reverse from year to year, depending on whether climatic variability across years is autocorrelated (positive or negative) or is uncorrelated/unpredictable environmental noise (Schwager et al. 2006; Ruokolainen et al. 2009). Offspring from perennial organisms produced during a current year may be the product of selection on both the present and past years. When the environment is positively autocorrelated, a given year closely reflects the following year’s environment. Such multi-year exposure can lead to stronger selection (Postuma *et al.,* 2020) due to a compounding effect of selection across multiple years. Thus, positive autocorrelation could increase the probability of rapid evolution and evolutionary rescue through climate change (Peniston *et al.,* 2020). However, positive temporal autocorrelation can also result in increased risk of extinction due to the compounded negative impact of multiple severe years if populations cannot adapt fast enough (Metcalf and Koons 2007; Ruokolainen et al. 2009). This could be particularly true during climate change where repeated exposure to extreme conditions over multiple years could push species closer to biological limits.

Whether current year or legacy effects are more important in describing rapid evolution may depend on the amount of standing genetic variation within a population, the timing of selection relative to demography, and the life history of the organism. Changes in the amount of standing genetic variation could make populations less able to rapidly evolve during certain years, especially given declining population sizes or population bottlenecks due to sequential climatic anomalies (Bell 2013; Pauls et al. 2013; Maron et al. 2015). In plants and other organisms with resting propagules (e.g., seed banks), individuals can recruit one, two, or more years after selection occurred on their parents (Brown and Venable 1986), making prior environments even more important to consider in determining the phenotype observed during a given year. Finally, life-history traits such as longevity and generation time are critically important to determining if, and when, evolution can occur. If a species has an annual lifecycle, then the observed year is likely to play a dominant role in selection (Friedman 2020). In perennial species the current year could still play a dominant role if selection mainly acts on reproductive plants. However, prior years may become more important (Blonder et al. 2023), especially if selection acts on juvenile plants that will not be reproductive until subsequent years. These effects of selection may not be seen as increased recruitment until years later because the selected individuals have not yet had time to reproduce. Thus, the effect of prior years may be vital in explaining rapid evolution in longer-lived species. Additionally, if the observed year also exerts selection on adult plants, then both prior and current years or the cumulative averages across multiple years may be most important in describing rapid evolution.

Drought and rising temperatures are two major stressors of climate change (Pörtner et al. 2022). These may act as a strong selective force driving rapid evolution but may also cause a large amount of cyclical variability across years, particularly in Mediterranean systems (Lionello et al. 2006; Deitch et al. 2017). Mediterranean systems are primarily known for having a strong influence of seasonal water availability, with a reduction in available water occurring during the mid-growing season, as well as multi-year cycles of drought (Deitch et al. 2017). As seen in a recent record-setting drought in California, the intensity and duration of these extreme weather events are increasing (Robeson 2015). Thus, the Mediterranean biome offers an excellent case study to examine the effects of climate change on evolution, given how global climate models predict increased drought with temperature increases of 1.5 to 2°C above pre-industrial levels (Xu et al. 2019).

Drought adaptation may occur through two main mechanisms in plants: dehydration avoidance and dehydration escape (Kooyers 2015; Volaire 2018). Avoidance occurs through morphological changes to prevent heat stress or water loss of the plant, such as closing stomata, having thicker leaves, and growing slowly. In contrast, during escape, plants speed up phenology to reproduce prior to the onset of more extreme and lethal conditions. The evolution of escape in response to drought has been widely documented in annual species (Franks et al. 2007; Ivey and Carr 2012; Paccard et al. 2014; Kooyers 2015), although in some cases avoidance appears to have evolved instead (Kooyers et al. 2019; Anstett et al. 2021). Dehydration avoidance might be more common in perennial species, which may benefit from avoidance across multiple years of growth and may be less able to escape drought due to greater longevity.

We recently found evidence of differential rapid evolution across the latitudinal range of *Mimulus cardinalis* over seven years that encompassed a record-setting drought (Anstett et al. 2021). Our results suggest evolution towards dehydration avoidance in central and southern parts of the range, whereby plants are flowering later and have smaller leaves (lower SLA) while little change occurred in the northern populations. We demonstrated change over time, but have not pinpointed which changes in climate and which time intervals and climatic parameters were likely exerting selection. This is important to investigate under different treatments, such as wet and dry conditions, because past climates could alter how populations respond to future stress. Here we assess different climate variables, Standardised Precipitation Evapotranspiration Index (SPEI), Hargreaves Climate Moisture Deficit Anomaly (CMDA), Mean Annual Precipitation Anomaly (MAPA), and Mean Annual Temperature Anomaly (MATA), during the aforementioned severe drought in California and Oregon.

We ask 1) what climatic variable best describes potential selective pressures in *Mimulus cardinalis*?; 2) is rapid evolution best described from legacies from past climates or the current climate (where climate is separated into 5 categories according to climate period: collection year, previous year, two-years previous, a cumulative effect of the current and previous year, and three-year cumulative) and how does this vary across regions and water treatments?; and 3) is strength of selection, and therefore rate of trait change, enhanced by positive temporal autocorrelation? Because selection is imposed by multiple aspects of climate, and the effects of drought are exacerbated by heat, we predict that a compound index like SPEI and CMDA should be a better predictor of trait evolution than MATA or MAPA. Prior years could be important in driving selection/evolution, because *M. cardinalis* is a perennial. Therefore, we predicted that short legacy effects encompassed by prior-year climate values and cumulative climate metrics from multiple years should outperform climate of the collection year in explaining trait evolution. We also expected legacy effects to be less evident in the southern part of the range where *M. cardinalis* has faster, more annualized life history compared to the slower and consistently perennial life history of the north. Finally, we predicted that positive temporal autocorrelation would increase the consistency of directional selection and yield greater amounts of trait change than in environments where the direction of selection is unpredictable or fluctuating.

## Methods

### Study System

We studied *Mimulus* (*Erythranthe*) *cardinalis* (scarlet monkeyflower), an herbaceous perennial plant that is found along riparian areas from central Oregon, USA to northern Baja California, Mexico. The species’ range includes areas affected by a recent drought of unprecedented severity that occurred in the western United States between the years 2012-2015 (Robeson 2015). Seeds were collected throughout the range of *M. cardinalis* prior to, during, and at the end of the drought. This gave a timeseries of seed collections from twelve populations from years 2010 through to 2016, representing three regions across the range: four populations from the north, five populations from the centre, and three populations from the south (Fig. 1A-C). These regions were determined by climatic differences and geographical continuity. Seeds from these collections were used for a resurrection experiment (Franks et al. 2018), whereby we compared ancestors and descendants in a greenhouse common garden to measure genetically-based phenotypic differences in drought adaptation. This was performed after first conducting a refresher generation (Franks et al. 2018) to reduce differences in seed age and maternal environment. Seeds were checked for the invisible fraction problem (see Anstett et al. (2021) for details). Most *M. cardinalis* individuals live 1-2 years with most individuals flowering for one or two growing seasons. Our seed collection timeseries encompasses 7 growing seasons, which is sufficient to observe turn-over in generations and therefore differences in phenotypes that can be attributed to genetic change as opposed to sampling idiosyncrasy of different subsets of the persistent individuals. While some individuals could have been sampled for more than one season, *M. cardinalis* is outcrossing and thus even in such an extreme case we would still be sampling a distinct gene pool. Thus, comparisons carried out in the greenhouse mostly reflect ancestor and descendant lineages across years with some amount of generational overlap.

**Figure 1.**
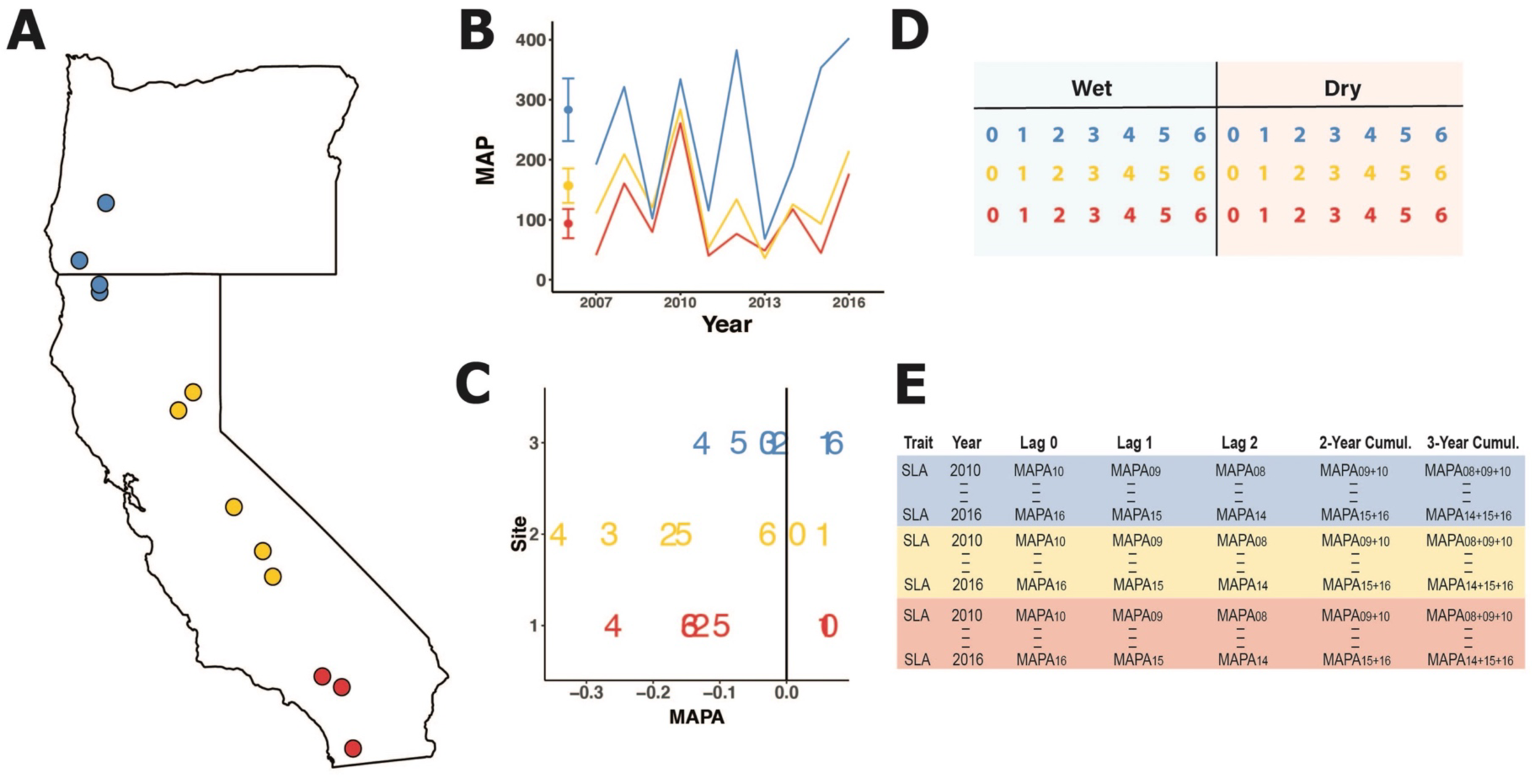
Schematic diagram depicting how the data were generated. A) Map of twelve focal populations of *Mimulus cardinalis* in Oregon and California, USA, where populations from the northern region are indicated in blue, centre region in yellow, and southern region in red. B) Mean annual precipitation (MAP) across 30-year average (dot and 95% CI) and MAP across time of interest in three selected populations representing each region. C) MAP anomaly (MAPA) calculated by subtracting the 30-year mean from annual MAP for the same three populations. Each number represents the last digit of the collection year (e.g., 2 = 2012). Negative values indicate years that were drier than average. D) Schematic of common garden greenhouse experiment illustrating how seeds from all sites and collection years were grown in wet and dry environments. There were multiple replicates for each family in a randomized block design (not shown). Numbers 0 to 6 reflect that genetic material from seeds collected from 2010 through to 2016 was included in the greenhouse experiment. E) Example of data frame configuration, showing trait, year of collection, and how each lag and cumulative measure was calculated.

### Growth Treatments and Data Collection

Seeds were sown on sand and misted until germinated (N = 3080 plants; see Anstett et al. (2021) for full design). Plant benches were flooded four times daily for 81 days until seedlings established. Once established, the plants were arranged in eight randomized blocks, four of which were assigned to a dry treatment and four assigned to a wet treatment (Fig. 1D). Position of the blocks alternated across greenhouse benches to account for variability in the greenhouse climate. All plants were initially watered four times a day until they were established. Treatments began on experiment day 82. This day was chosen to reflect the plant life stage at which natural drought impacts can become considerable in the field. Plants in the wet treatment were given the same amount of water they had been receiving up until 81 days (watering four times a day = 100%). The dry treatment was a dry down established by reducing watering by 50% on day 82 and by another 50% on day 96. Dry treatment plants were no longer watered on day 111. This set up was used to mimic natural drying during the summer growing season of a Mediterranean climate. Plants were monitored daily to record the date of first flowering. In three out of the four blocks (3 replicates) we sampled one leaf from each plant, removing the leaf from the sixth to eighth node for specific leaf area (SLA; area/dry leaf mass) measurements. Leaves were removed 15-22 days following the beginning of the dry treatment to ensure leaves had developed under the conditions of interest. Because of the refresher generation and randomized common garden design, all phenotypic differences exhibited are attributable to evolved genetic differences, apart from possible transgenerational plasticity and potential rare repeated collections of perennial individuals.

### Climate Data

To determine which climatic variable best described potential selective pressures (question 1), we examined four climate parameters: standardised precipitation evapotranspiration index (SPEI; negative refers to drought), Hargreaves climate moisture deficit anomaly (CMDA; positive refers to drought), log mean annual precipitation anomaly (MAPA; negative refers to drought), and mean annual temperature anomaly (MATA; positive refers to higher temperature). SPEI was downloaded from the Global SPEI Database at (spei.csic.es/database.html) and was calculated for each 0.5-degree grid closest to each site for the water year (October of previous year to September of collection year, which is biologically relevant for *M. cardinalis*, which senesces in September) (Fig. S1). Climate moisture deficit (CMD), mean annual precipitation (MAP), and mean annual temperature (MAT) were downloaded as monthly data from Climate NA for each population (Wang et al. 2010). We then used monthly data to calculate water year from 1980 to 2016 (October of previous year to September of given year). A 30-year average was also calculated from October 1979 to September 2009. Annual anomalies for each climate variable during the sampling period were calculated as outlined in Anstett et al. (2021) (Fig. S2-S4). Briefly anomaly = yearly value −30yr average. Both SPEI and CMD are drought indices. CMD examines a finer scale of site-level differences in evaporation and precipitation integrated across months, whereas SPEI examines drought at a broader regional scale, but incorporates additional metrics of hydrology.

To understand which climate period best describes the observed trait change (question 2), we evaluated three single-year periods: the collection year (lag 0), the year previous (lag 1), and two-years prior (lag 2). We also evaluated cumulative effect of the current and previous year (2-year lag) by summing lag 0 and lag1, and the cumulative effect of the collection year and the two years prior (3-year lag) by summing lag 0, lag 1 and lag 2 (Fig. 1E).

### Statistical Analysis

To test for explanatory power of the different climate variables between models (question 1) and different time/lag periods within each climate variable (question 2) we employed a model selection approach using AIC (Akaike 1974; Findlay and Bourdages 2000). We analyzed each region by treatment (i.e., north-wet and north-dry) combination separately focusing on one climate-year variable at a time (e.g., MAPA lag 0). We used an additive mixed effects model with block and climate-year as fixed effects, and year and full-sibling family nested within each site as random effects. Year was included as a random effect to account for additional differences between years that are not indexed by the climate variables. Models were run with the *lmer* function in the lme4 package (Bates et al. 2014) for R in version 4.0.4. (R Development Core Team 2020) separately for SLA and date of flowering and for each lag and cumulative metric across SPEI, CMDA, MAPA, and MATA. We were unable to use different climate variables within the same model because of collinearity (Table S1). We were also unable to use multiple lags in the same model within climate variables because of collinearity, especially for MATA which had Pearson r = ∼0.7 between lags and averages.

We compared across all models within each region-year combination to determine the best climate variables and periods through AIC comparison (delta AIC < 2). The AIC values were calculated based on maximum likelihood using the AIC function in R. We did not have any missing data for the climate-year or block, making our models comparable. We further assessed these models through likelihood ratio tests on nested models using the *lrtest* command from the lmtest package (Hothorn et al. 2019) to assess statistical support for model terms. This involved comparing each AIC selected model with one climate lag to a model with only block and random effects. For each trait, P-values were corrected for multiple testing using a Bonferroni correction within region and treatment. Marginal and conditional R-squared values were assessed using the function *r.squaredGLMM* in the MuMIn package (Barton and Barton 2019). We extracted the model predictions for the factor of interest for all favoured models using *visreg* (Breheny et al. 2020) and plotted these residuals, grouping them appropriately according to the selected model, using *ggplot2* (Wickham 2011).

The analyses above identify which aspects of climate, over which climate periods, best predict trait differences, but they remove information about the sequence of climate conditions. To test whether sequences of similar climate anomalies lead to more trait change than more variable climate (question 4), we modeled trait change as a function of autocorrelation with the prior year’s value (lag 1) in climate across the timeseries (2010-2016) within wet and dry treatments using *acf* function in the tseries package (Trapletti et al. 2015). The metric varies as a −1 to +1 continuum from negative to positive autocorrelation. For each of the 12 sites, we expressed trait change as the absolute value of the trait at the end of the timeseries minus the trait at the beginning of the timeseries. Absolute trait change was considered because we were not interested in the direction of change, but rather the amount of trait change. We included an interaction between autocorrelation and mean climate anomaly to ensure the model could treat autocorrelation across benign periods differently to autocorrelation across extremely anomalous periods. In other words, selection imposed by a series of benign years should be much weaker than selection imposed by a series of anomalous years.

## Results

The best climatic variable (delta AIC < 2) for explaining the evolution of both SLA and date of flowering was mean annual precipitation anomaly (MAPA), when comparing across all climate variables (MAPA, CMDA, MATA, SPEI) within each region (north, centre, and south) and treatment (wet and dry; Table S2). The one exception to this was MATA, which was also a favoured explanatory variable in the south under the dry treatments, but only for the evolution of SLA. The climate periods that best explained rapid evolution of SLA and flowering time differed between regions and across treatments. While current-year climate (lag 0) was present in many of the selected models, it was not always favoured. Geography and treatment affected whether lag 0 was favored and if it was either associated with similar or conflicting patterns for trait evolution. All AIC selected linear mixed models were significant after a Bonferroni correction, with the exception of models for center wet and south dry treatments. Marginal R-squared ranged from 0.02 – 0.27 (Table 1).

**Table 1.** P-value and R^2^ of climate lags and cumulative metric explaining evolution in Specific Leaf Area (SLA) and Flowering Time (FT). All models have the climate-year (for a given environmental variable) and block as additive fixed effects, with year and full-sibling family nested within each site as random effects. Each model is tested against a simpler model with the climate variable removed using likelihood ratio tests. Delta loglik (between the two models), p-value, marginal R^2^ and conditional R^2^ are given for each model. SPEI = Standardised Precipitation Evapotranspiration Index; CMDA = Hargreaves Climate Moisture Deficit Anomaly; MAPA = log Mean Annual Precipitation Anomaly; MATA = Mean Annual Temperature Anomaly. Lag 0 = effect of current year’s climate; lag 1 = effect of climate from one year prior, lag 2 = effect of climate from two years prior; 2-year = cumulative effect of lag 0 and lag 1; 3-year = cumulative effect of lag 0, lag 1, and lag 2. Bolding is given for significant p-values after a Bonferroni correction carried out within each region, treatment, and trait. Underlined is given for marginally significant after a Bonferroni correction.

| Region | Treatment | Trait | Environmental Variable | Lag | Delta LogLik | P-value | Marginal R<sup>2</sup> | Conditional R<sup>2</sup> |
| --- | --- | --- | --- | --- | --- | --- | --- | --- |
| North | Wet | SLA | MAPA | lag0 | 7.38 | **0.0001** | 0.11 | 0.18 |
|  |  | FT | MAPA | lag0 | 3.36 | **0.01** | 0.04 | 0.21 |
|  |  | FT | MAPA | lag1 | 3.02 | **0.014** | 0.04 | 0.20 |
|  |  | FT | MAPA | lag2 | 3.93 | **0.005** | 0.05 | 0.21 |
| North | Dry | SLA | MAPA | lag0 | 7.90 | **0.0001** | 0.27 | 0.34 |
|  |  | FT | MAPA | lag0 | 3.96 | **0.005** | 0.06 | 0.22 |
|  |  | FT | MAPA | lag1 | 3.43 | **0.009** | 0.06 | 0.22 |
|  |  | FT | MAPA | lag2 | 3.50 | **0.008** | 0.06 | 0.24 |
|  |  | FT | MAPA | 2-year | 4.14 | **0.004** | 0.06 | 0.22 |
|  |  | FT | MAPA | 3-year | 3.39 | **0.009** | 0.06 | 0.22 |
| Centre | Wet | SLA | MAPA | lag0 | 4.48 | **0.003** | 0.09 | 0.16 |
|  |  | SLA | MAPA | lag1 | 4.49 | **0.003** | 0.09 | 0.17 |
|  |  | SLA | MAPA | lag2 | 5.46 | **0.001** | 0.10 | 0.17 |
|  |  | SLA | MAPA | 3-year | 4.80 | **0.002** | 0.10 | 0.17 |
|  |  | FT | MAPA | lag0 | 2.81 | <u>0.018</u> | 0.07 | 0.21 |
|  |  | FT | MAPA | lag1 | 2.50 | 0.025 | 0.07 | 0.21 |
|  |  | FT | MAPA | lag2 | 2.83 | <u>0.017</u> | 0.07 | 0.21 |
|  |  | FT | MAPA | 2-year | 2.23 | 0.035 | 0.07 | 0.21 |
|  |  | FT | MAPA | 3-year | 2.39 | 0.029 | 0.07 | 0.21 |
| **Centre** | **Dry** | SLA | MAPA | lag0 | 4.90 | **0.002** | 0.20 | 0.26 |
|  |  | SLA | MAPA | lag1 | 5.42 | **0.001** | 0.20 | 0.26 |
|  |  | SLA | MAPA | lag2 | 4.72 | **0.002** | 0.19 | 0.26 |
|  |  | FT | MAPA | lag0 | 3.10 | **0.013** | 0.07 | 0.30 |
|  |  | FT | MAPA | lag2 | 3.50 | **0.008** | 0.07 | 0.30 |
| **South** | **Wet** | SLA | MAPA | lag1 | 6.59 | **0.0003** | 0.11 | 0.27 |
|  |  | SLA | MAPA | lag2 | 6.28 | **0.0004** | 0.11 | 0.28 |
|  |  | FT | MAPA | lag2 | 4.92 | **0.002** | 0.11 | 0.48 |
|  |  | FT | MAPA | 3-year | 4.78 | **0.002** | 0.18 | 0.56 |
| **South** | **Dry** | SLA | MAPA | lag0 | 5.84 | **0.001** | 0.23 | 0.34 |
|  |  | SLA | MAPA | lag1 | 5.00 | **0.002** | 0.22 | 0.34 |
|  |  | SLA | MAPA | lag2 | 4.88 | **0.002** | 0.22 | 0.34 |
|  |  | SLA | MAPA | 2-year | 5.11 | **0.001** | 0.23 | 0.34 |
|  |  | SLA | MATA | lag0 | 5.72 | **0.001** | 0.25 | 0.33 |
|  |  | FT | MAPA | lag0 | 2.29 | 0.032 | 0.02 | 0.31 |
|  |  | FT | MAPA | lag1 | 2.40 | 0.028 | 0.02 | 0.30 |
|  |  | FT | MAPA | lag2 | 2.41 | 0.028 | 0.02 | 0.30 |
|  |  | FT | MAPA | 2-year | 1.73 | 0.063 | 0.02 | 0.31 |
|  |  | FT | MAPA | 3-year | 1.62 | 0.072 | 0.02 | 0.30 |

### North

For SLA, lag 0 was favoured under wet treatments (Fig. 2A) and a broader leaf (greater SLA) was associated with decreased MAPA (drier conditions) of the current year. In dry treatments, lag 2 was favoured and showed the opposite trend, broader leaf with increased MAPA from 2 years prior (Fig. 2B). For flowering date, all single-year lags (0, 1, and 2) were favoured in wet treatments and all lags and averages were favoured in dry treatments (Fig 2, Fig S5). In both treatments later flowering was associated with decreased MAPA in the current year (lag 0; Fig. 2C,E), while later flowering was associated with increased MAPA of 2 years prior (Fig. 2D,F). Additionally, Lag 1 showed a weak association with date of flowering in the wet treatment (Fig. S5A) and all other dry lags showed the same pattern as lag 0 in the dry treatment (Fig. S5B-D).

**Figure 2.**
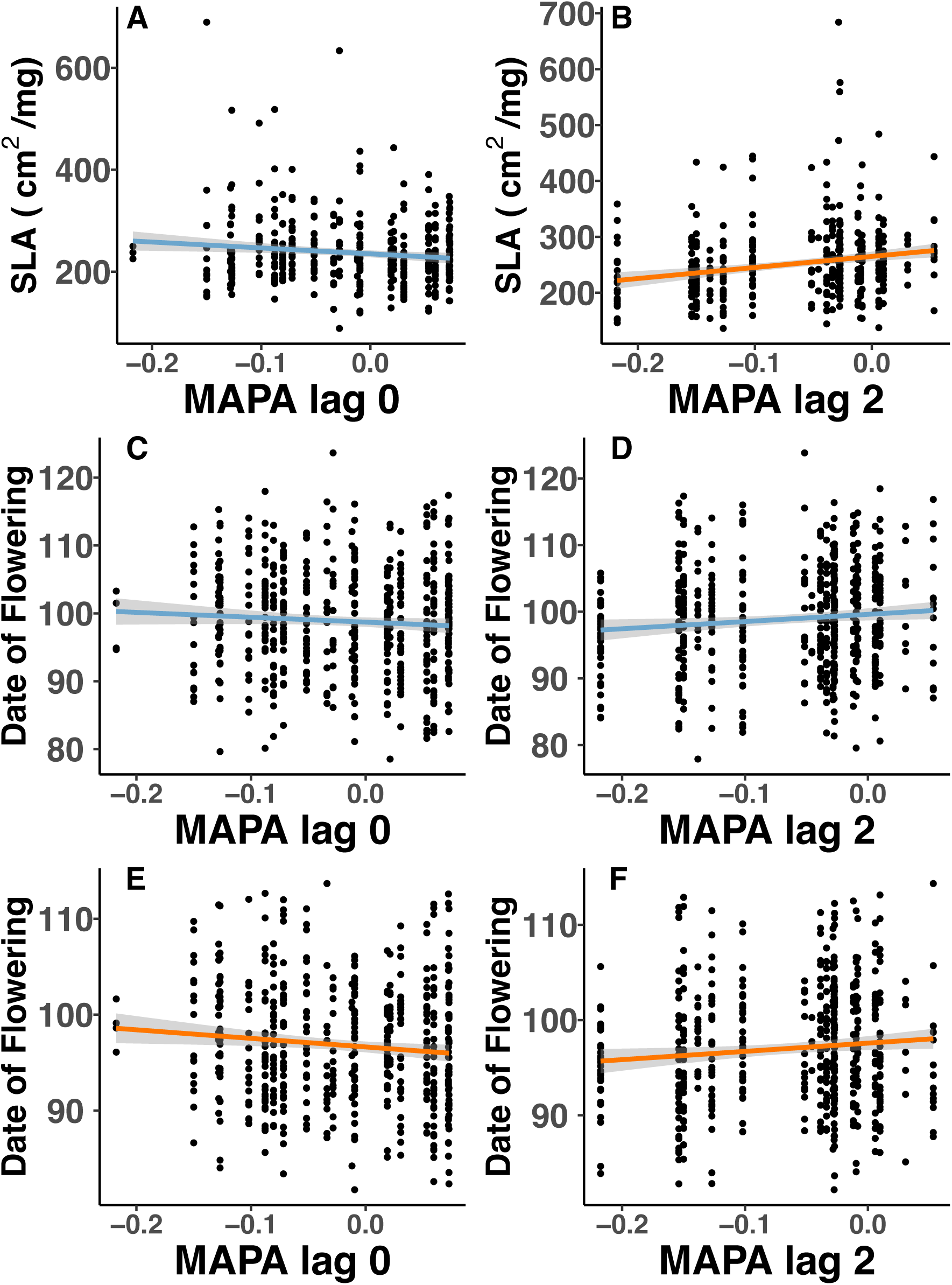
Evolution of Specific Leaf Area (SLA) and date of flowering across mean annual precipitation anomaly (MAPA) in the northern part of the *Mimulus cardinalis* range. A) SLA is best explained by lag 0 when plants are grown under a well-watered treatment. B) SLA is best explained by lag 2 when plants grown under a drought treatment. C) Date of flowering is explained by lag 0 and D) lag 2 when grown under a well-watered treatment. E) Date of flowering is explained by lag 0 and F) lag 2 when plants grown under a drought treatment. Lag 0 = effect of current year’s climate; lag 2 = effect of climate from two years prior. Blue regression line = well-watered treatment; orange regression line = drought treatment.

### Centre

In wet treatments, all lags except 2-year cumulative lag were favoured for SLA. All models showed the same pattern, having slightly thicker and narrower leaves when drought had occurred in the prior years (Fig. 3A,B; Fig. S6A,B). Under dry treatments, all single-year lags were favoured (0, 1, 2) and patterns were conflicted. Lag 0 had thicker leaves when the current year experienced drought (Fig. 3C), while lag 1 and 2 had broader, thinner leaves when one of the prior two years were under drought (Fig. 3D; Fig. S6C). For flowering time in wet treatments, all climate lags were explanatory and showed a general trend of earlier flowering time when drought had occurred earlier (Fig. 3E, Fig. S6D-F), except for lag 1, which showed no change (Fig. 3F). Under dry treatments, only lag 0 and 2 were favoured for flowering time in the centre under dry treatments and each showed opposite patterns. Later flowering occurred when the current year experienced drought (Fig 3G), while earlier flowering was observed when drought occurred two-years prior (Fig 3H).

**Figure 3.**
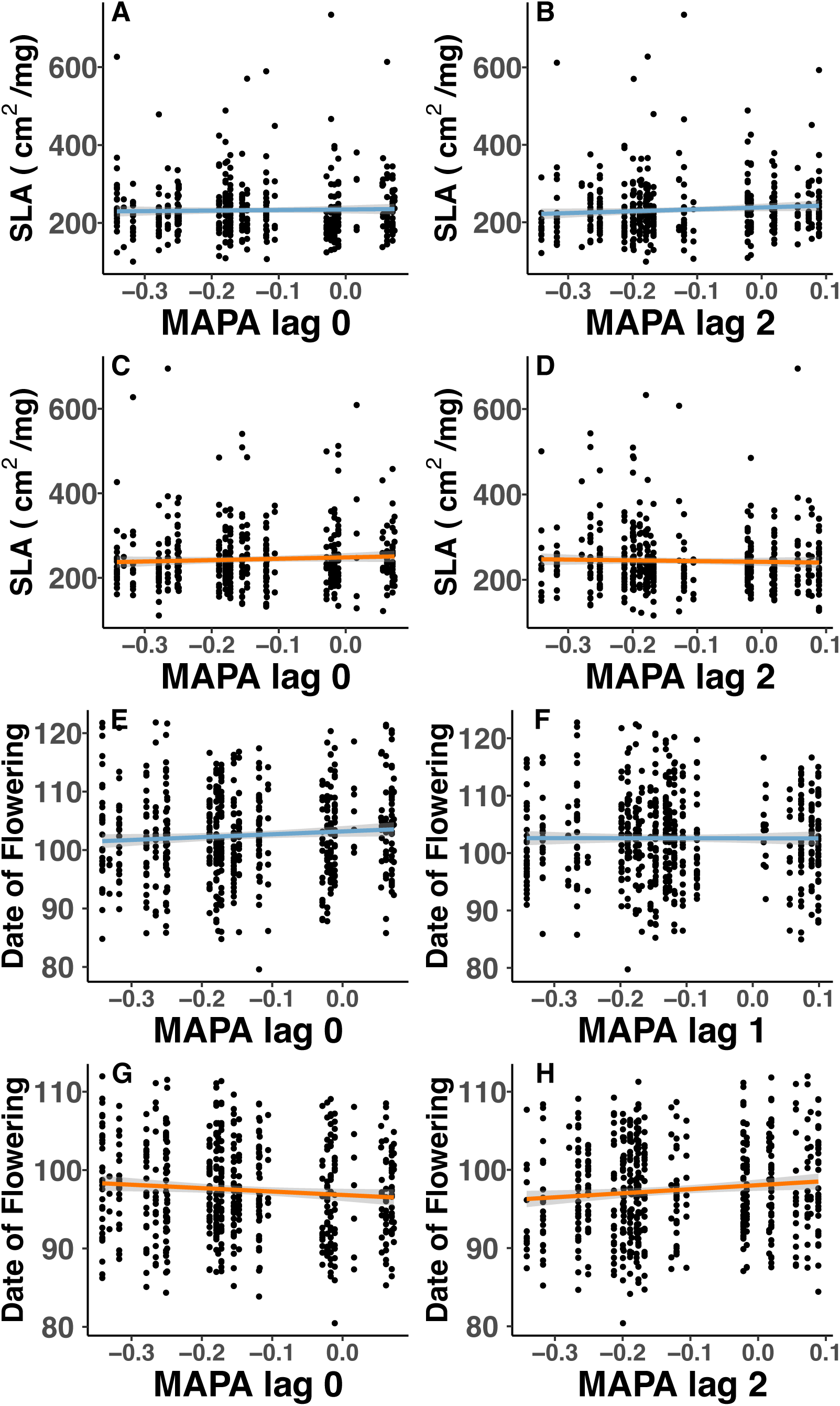
Evolution of Specific Leaf Area (SLA) and date of flowering across mean annual precipitation anomaly (MAPA) in the central part of the *Mimulus cardinalis* range. A) SLA is explained by lag 0 and B) lag 2 when grown under a well-watered treatment. C) SLA is explained by lag 0 and D) lag 2 when plants grown under a drought treatment. E) Date of flowering is explained by lag 0 and F) lag 1 when grown under a well-watered treatment. G) Date of flowering is explained by lag 0 and H) lag 2 when plants grown under a drought treatment. Lag 0 = effect of current year’s climate; lag 1 = effect of climate from one year prior; lag 2 = effect of climate from two years prior. Blue regression line = well-watered treatment; orange regression line = drought treatment.

### South

For SLA in wet treatments, lag 1 and 2 were favoured with broader leaves associated with lower MAPA (Fig. 4A; Fig. S7A). In dry treatments, MATA (Mean Annual Temperature Anomaly) lag 0 was favoured with thicker leaves when the current year was hotter (Fig. 4B). Additionally, in the dry treatment for SLA, all MAPA models except for 3-year cumulative were favoured. All of these lags showed thicker and smaller leaves following drought years (Fig. 4C; Fig. S7B,C) except for lag 2 which showed no pattern (Fig. 4D). For flowering time in wet treatments, lag 2 and the 3-year cumulative models were favoured with a strong negative association between MAPA and date of flowering (Fig. 4E; Fig S7D). All models were included for date of flowering under dry treatments but no trends were clearly observed (Fig. 4F-H, Fig. S7EF).

**Figure 4.**
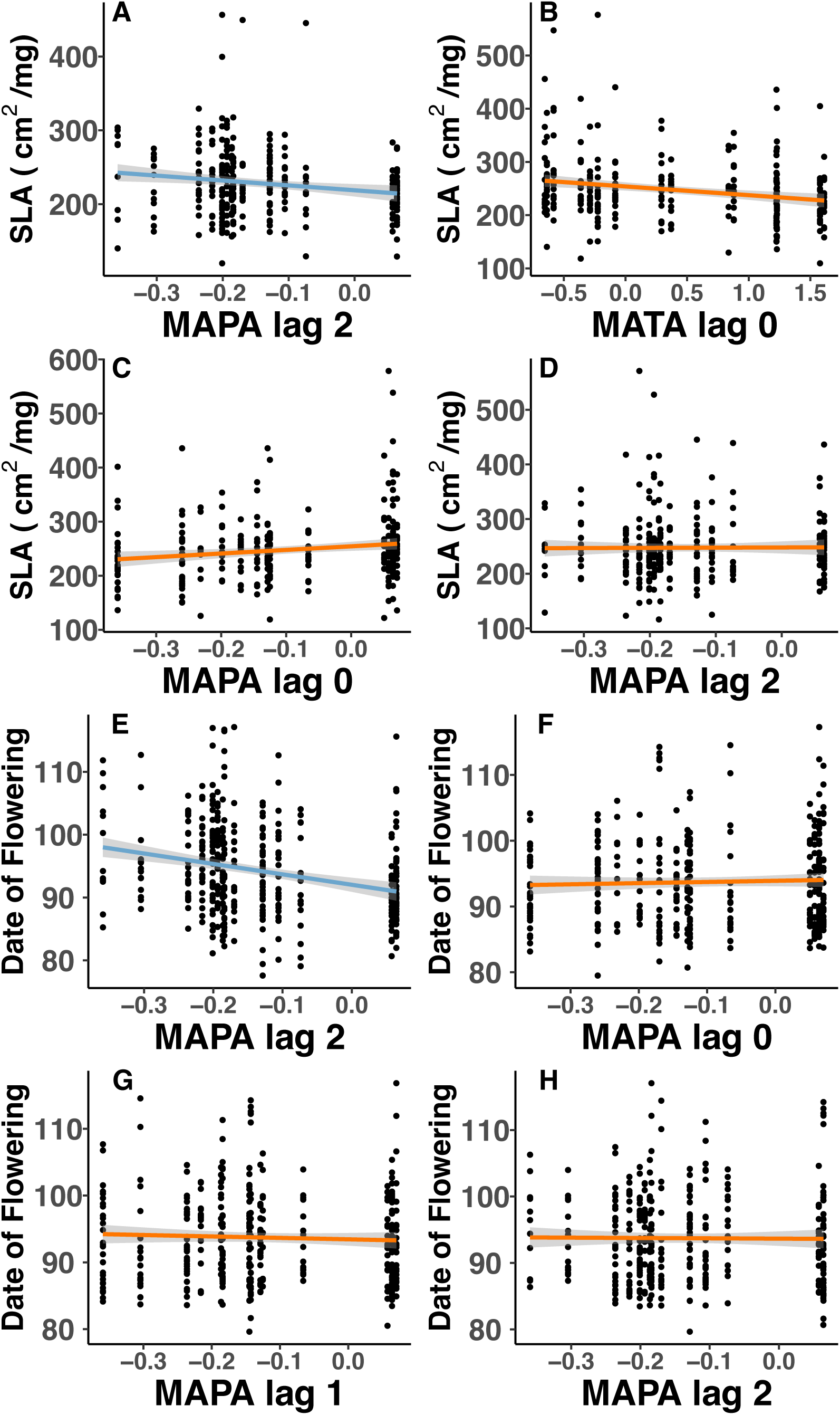
Evolution of Specific Leaf Area (SLA) and date of flowering across mean annual precipitation anomaly (MAPA) and mean annual temperature anomaly (MATA) in the southern part of the *Mimulus cardinalis* range. A) SLA is explained by MAPA lag 2 when grown under a well-watered treatment. SLA is also explained by B) MATA lag 0, C) MAPA lag 0 and D) MAPA lag 2) when plants grown under a drought treatment. E) Date of flowering is explained by MAPA lag 2 when grown under a well-watered treatment. Date of flowering is also explained by F) MAPA lag 0, G) MAPA lag 1, and H) MAPA lag 2 when grown under a drought treatment. Lag 0 = effect of current year’s climate; lag 1 = effect of climate from one year prior; lag 2 = effect of climate from two years prior. Blue regression line = well-watered treatment; orange regression line = drought treatment.

### Interannual climate

We found little support that positive autocorrelation affected the degree of trait change (Fig. S8-S9, Table S3). There was weak support that interannual MAPA had an effect on the rate of change of SLA under dry treatments (p = 0.086, R^2^ = 0.25; Fig. S4D, Table S3), showing greater change when MAPA was autocorrelated across years and less trait change when interannual climate was less predictable. When the climate time series was divided into subsets, we still found little evidence for a relationship between climate autocorrelation and trait change (Table S4; Fig. S10-S11).

## Discussion

Here we examined rapid evolution of two key traits related to drought avoidance and escape in relation to anomalies in four climatic variables across different time windows. Mean annual precipitation anomaly (MAPA) best explained patterns of rapid evolution compared to anomalies in temperature or drought indices. This was true except for southern plants under a dry treatment, where mean annual temperature anomaly (MATA) was also selected. There were regional and treatment differences in the importance of lags and cumulative time periods. When the climates of both current and previous years were selected as explanatory models, there was often conflict in the trait direction. While we did not detect a relationship between autocorrelation and trait evolution, we did find that phenotypic lags associated with past years (one-to two-years prior) often explained the direction of trait evolution as well or better than the current environment. For instance, a lag of one or two years might show an opposite pattern compared to lag 0. This suggests that the outcomes of rapid evolution across years observed in Anstett et al. (2021) are frequently the end result of conflicting selection across present and past years. Legacy effects of past climates on phenotypes are well documented in the ecological literature when studying phenomena such as community-level phenology, masting, and demographic vital rates (Schauber et al. 2002; Parmenter et al. 2018; Ascoli et al. 2020; Evers et al. 2021). Here we extend this type of inference and considered the impacts of phenotypic lags on the evolution of phenology and leaf morphology within a perennial species. Our findings suggest that predictions and explanations for how populations respond to specific climatic events might be less accurate if they do not consider the lagging effects of past climates.

### Importance of Previous Climates for Interpreting Evolutionary Responses

It is widely acknowledged that trait values observed in the present-day have been shaped by past selection (Schwager et al. 2006; Metcalf and Koons 2007; Fischer et al. 2011; Wieczynski et al. 2018) and yet it is unclear how much weight should be placed on past climates when studying rapid evolution. Generally, studies seeking to relate rapid phenotypic change to climatic change do so by studying the effect of a given year’s environment, the environments in which those phenotypes were expressed (i.e. lag 0), on trait change (Franks et al. 2007; Bell and Gonzalez 2011; Anstett et al. 2021). We previously found that rapid evolution in SLA and flowering time varied across different regions of *M. cardinalis,* with greater evolutionary changes in populations from central and southern regions (Anstett et al. 2021). This analysis occurred from a pre-drought condition up to a peak drought year, inferring that decreased water availability and increased temperature drove these patterns, but without specifically including climate variables as predictors.

Here we found that climate data from past years compared to the collection year often resulted in different interpretations for the way that populations responded to environmental change. For example, interpretations of northern populations, made by only considering the collection year (lag 0) for mean annual precipitation anomaly (MAPA), suggested evolution of later flowering in dry years and earlier flowering in wet years. However, the effect of MAPA from two-years-prior (lag 2) suggested evolution of earlier flowering in dry years and later flowering in wet years: the exact opposite pattern as lag 0 (wet: Fig 2C,D; dry: Fig. 2E,F). This alternating selection likely explains one of our main findings in Anstett *et al*. (2021): lack of evolutionary change in the northern populations of *M. cardinalis* for SLA and relatively weak change for flowering time. These findings are likely due to legacy effects on selection across climatic conditions from prior years and these patterns were not directly revealed by regressing trait values against year of collection or by merely including climate from year of collection.

The conflicting trait directionality in the northern populations could be due to alternating selection caused by shorter drought duration and intensity in the Northern range (Fig. S3). A shorter drought period would likely result in prior years selecting for different phenotypes than current years (e.g., lag 0 = wetter, lag 2 = drier). Another reason for the importance of two-years prior for the northern populations could be the perennial nature of *M. cardinalis*, which is more prominent in the North. Selection might have occurred on parental phenotypes selecting for leaf traits or phenology that are not optimal for the current year’s climate; however, the impact of this selection might be most clearly seen two-years later when the surviving individuals might be contributing the most to the gene pool. Alternatively, the relative importance of lags in the northern populations could be due to non-genetic inheritable traits, for instance transgenerational plasticity. Transgenerational plasticity has been observed to play a role in *M. cardinalis* phenotype (Branch, 2023). The lasting effects from previous generations are becoming more well-documented in plants, showing that environments of grandparents (lag 2) can alter phenotypes of grand-offspring (Groot et al. 2016; Rajpal et al. 2022), and are not fully removed by using refresher generations. Thus our findings could also provide evidence of the importance of epigenetic changes affecting evolution in the northern populations of *M. cardinalis*.

Similar to our findings, Mulder *et. al.* (2017) found the timing of flowering across species was associated with the temperature from one-or two-years prior in perennials. Their study highlighted different responses across species, where one-year lags were observed in earlier flowering species and current-year and two-year lags were observed in later flowering species. Contradictory patterns were also observed in their study, where warmer temperatures of the current year caused earlier flowering in some species, but warmer temperatures from two-years prior caused later flowering (Mulder et al. 2017). While they did not examine evolutionary consequences of these temperature changes, their community-level results corroborate the effects observed in our study. These results further suggest that climatic variability may reduce expected evolutionary change, even on a short timescale.

Overall, we found that mean precipitation anomaly often explained evolutionary change, although the temporal scale and directionality of this legacy varied across regions and treatments. In contrast to northern populations, the impact of precipitation and temperature on trait change for central and southern regions found in this study largely retained the same pattern across climate lags. Additionally, these patterns generally corroborated the observed changes over time seen in Anstett et al. (2021). For example, SLA decreased in dry treatments for southern genotypes from pre to peak drought year (Anstett et al. 2021), which we also observed for southern genotypes under dry climatic anomalies in this study (Fig. 4C; Fig. S7B,C). This decrease occurred for lag 0 (Fig. 4C), lag 1 (Fig. S7B), and 2-year cumulative models (Fig. S7C). For SLA under wet treatments in the southern region, only prior years (lag 1 and 2) were selected and predictive of the results seen in Anstett *et al*. (2021), showing that SLA increased following dry years. Similarly, lag 2 and 3-year cumulative models were selected for flowering date under wet treatments in the southern populations, showing later flowering following dry years. These findings show an unexpected importance of prior year climate as a driver of rapid evolution in Southern regions of the *M. cardinalis* range.

We expected that lagged effects would be less predictive in more arid Southern regions due to increased annualization; however, lagged effects commonly explained rapid evolution. There are two possible explanations for this discrepancy. Selection from prior years on perennial parental genotypes may be particularly strong in more arid regions due to larger mortality during non-reproductive years (Harrington 1991; Lewandrowski et al. 2021). Thus prior year selection might be more important than selection in the present year due to a small but well-adapted number of individuals that survived more than one season. In certain years, such impacts might also be exaggerated by pulsed recruitment events that are common within arid systems (Reynolds et al. 2004; Kenny and Moxham 2022). Alternatively, the impact of a seedbank may also produce a similar effect, where climatic impacts from two-years-prior might affect the phenotypes present in the current year due to prior years’ selection on the composition of the available seed bank. Seed banking is a common strategy to overcome climatic variability (Mayfield et al. 2014; Torres-Martínez et al. 2017), which results in a population of plants that reflects selection from prior climates. Little is known about seed banks in *M. cardinalis*, as they are difficult to study in unstable, sandy soils that characterize riparian systems. However, given that *M. cardinalis* becomes increasingly annualized in southern environments, a seed banking explanation would be congruent with the importance of lagged and multi-year impacts on trait evolution.

Overall, the lack of broad and consistent explanatory power of climate from the current year (lag 0) highlights the need for a broader consideration of the impacts of prior years when studying rapid evolution, particularly in non-annual species. Our results highlight how environments of different years can translate to different interpretations of results and can strengthen our understanding of other observed relationships (such as is seen for phenotypic lags in northern populations). These relationships show the importance of considering the impact of previous years when studying the evolutionary impact of climate.

### Importance of Different Climate Variables for Evolution

Researchers generally select climate variables *a priori* when analysing evolutionary data (Evers *et al*., 2021). We predicted that metrics for drought, such as SPEI and CMDA, would have the greatest explanatory power for trait change in our system, because they integrate precipitation and temperature, as well as evapotranspiration (Wang et al. 2012; Beguería et al. 2014). However, contrary to our prediction, MAPA was the best explanatory variable for trait evolution (Table S2). Overall, while precipitation was selected across regions throughout a drought cycle, it does not preclude drought metrics or temperature being more important within shorter timescales (e.g. a pre-drought to peak drought gradient; (Anstett et al. 2021)) or across multidecadal datasets. Indeed, these other metrics could be better at integrating the effect of climatic change on temperature and how it interacts with water availability, which may be more important during a progression from higher water availability to severe drought or across a longer timescale, where temperature changes may be more influential.

### Autocorrelation of Interannual Climate Variability

We predicted that positive climatic autocorrelation during an exceptionally severe drought would lead to higher rates of trait change due to an increase in selection consistency (Ruokolainen et al. 2009; Michel et al. 2014; Wieczynski et al. 2018; Postuma et al. 2020), since climatic autocorrelation has been associated with increased rate of adaptation (Peischl and Kirkpatrick 2012). However, we found no clear effect of climatic autocorrelation on trait evolution across the range of *M. cardinalis* (Table S3-S4; Fig. S5-S8). This could be due to low resolution to detect these patterns because we only examined seven years and not a broader temporal dataset, such as a 20-to 30-year timeline where these trends might become apparent (Burthe et al. 2016; Petriccione and Bricca 2019). It could also indicate that autocorrelation has not had a strong effect on rapid evolution in this system yet, although positive autocorrelation and hence selection pressure could increase as climate change intensifies.

Positive autocorrelation across years implies that consecutive years are likely to have a similar climate and thus higher predictability across years. However, variability within a season (e.g. increased heat waves and cool periods, periods of particularly low water availability followed by a pulse of higher water availability) are not captured by a yearly autocorrelation value, and may prove to be more important in influencing selection since an organism experiences the variation while it is alive. For example, the lower-than-predicted trait change of an alpine butterfly (*Colias meadii*) over 60 years of increased annual temperature was attributed to fluctuating selection from the additional increase in temperature variability that occurred during the same time (Kingsolver and Buckley 2015). In this way, increased variability, which decreases the rate of evolution, could counteract positive autocorrelation pressure that would otherwise increase the evolutionary rate and therefore decrease the impact of autocorrelation on selection from year-to-year.

### Caveats

One concern about associations between traits and climate in our system is the variability presented. Our model comparisons are all relative and visualizations reveal a large amount of scatter. There are two main explanations for this variability. First, the climate data are based on downscaled interpolations of PRISM data in a topographically complex riparian landscape (Wang et al. 2010; Wang et al. 2016), where microclimates and local-scale variability are considerable. Yet yearly flooding and the dynamic nature of riparian habitats make it largely unfeasible to consistently place and retrieve climate data loggers. Secondly, additional unmeasured climatic factors and biotic interactions may also be driving some of this variability. Overall, we present the best possible climate data available for this system. Even though the models have a large amount of scatter, we are still able to recover nuanced comparisons that are important to our understanding of both *M. cardinalis* and on how lagged effects may operate in water limited environments.

A separate concern is that we collected a single fruit per sampled plant during each year (2010-2016). Therefore, we equally weighted the gene pool of our refresher generation for each plant we sampled regardless if that plant had one or many fruits. Thus, our results likely underrepresent differential ability to produce additional progeny in plants that had greater fecundity. This type of bias makes it less likely for us to observe selection, a flaw with most resurrection experiments. Despite this bias, we are still able to observe rapid evolution across a small number of generations (Anstett et al. 2021), and across lags. Thus, this issue does not seem to diminish our ability to detect phenotypic change due to evolution.

## Conclusion

Overall, we show the importance of considering phenotypic lags as legacies from selection in past climates when investigating rapid evolution to climate change. These lag effects can be more important in explaining trait change than the effects of the current environment, particularly in longer-lived and seed banking species. Additionally, past time periods are important across multiple climate variables and lead to conflicting selection patterns on phenology and leaf traits. Further uncovering these phenotypic legacies is of considerable importance to better predict and interpret the impacts of future climate change. This is especially true in fluctuating environments, where phenotypic lags might lead to slower adaptation during extreme climate anomalies.

## Contributions

DA and AA developed the experiment. HB and DA collected the data, ran the analyses, and interpreted the results. HB and DA wrote the manuscript with edits by AA.

## Supporting information

https://www.dropbox.com/scl/fi/g6lx3ho8ltxqzydvcxx1i/Supplemental_Material_Branch_etal_bioRxiv.pdf?rlkey=6ndyfwd3i2h1tdilc2820l26b&dl=0

## Acknowledgements

A. Doherty, A. Wilkinson, M. Zink Yi, J. Zajonc, and K. Saller provided greenhouse support management and data collection. S. M. Robson provided advice on drought indices. J. Anstett, D. Turner and S. M. Robson provided R support. Special thanks to the Angert Lab 2022 for comments on an early version of the manuscript. Funding was provided by NSERC Banting, NSERC PDF, Killam, and Plant Resilience Institute Fellowships to D. Anstett. Funding was also provided by an NSERC PGS-D and Rick Hansen Man in Motion Fellowship to H. Branch, and NSERC Discovery grant to A. Angert.

## Data Accessibility Statement

The data and code are available on GitHub at https://github.com/anstettd/climate_lag

