## Supplementary material for "Phenotypic lags influence rapid evolution throughout a drought cycle": https://www.dropbox.com/scl/fi/g6lx3ho8ltxqzydvcxx1i/Supplemental_Material_Branch_etal_bioRxiv.pdf?rlkey=6ndyfwd3i2h1tdilc2820l26b&dl=0

**Supplemental Material for Branch et al.**

**Table S1.** Pearson correlation coefficients for four environmental variables across our timeseries (2010-2016) and our 12 sites with each site-year combination as the unit of replication.

| | SPEI | CMDA | MAPA | MATA |
|---|---|---|---|---|
| **SPEI** | 1 | -0.6 | -0.87 | -0.65 |
| **CMDA** | -0.6 | 1 | 0.45 | 0.77 |
| **MAPA** | -0.87 | 0.45 | 1 | 0.43 |
| **MATA** | -0.65 | 0.77 | 0.43 | 1 |

**Table S2.** Delta AIC values for range-wide models for specific leaf area (SLA) and date of flowering. A mixed effects model is run in all cases with Climate + Block as fixed effects. The best models (delta AIC<2) are indicated with asterisks. Within each climate variable, the best climate periods are indicated in bold. SPEI = Standardised Precipitation Evapotranspiration Index; CMDA = Hargreaves Climate Moisture Deficit Anomaly; MAPA = log Mean Annual Precipitation Anomaly; MATA = Mean Annual Temperature Anomaly. Lag 0 = effect of current year's climate; lag 1 = effect of climate from one year prior, lag 2 = effect of climate from two years prior; 2-year = cumulative effect of lag 0 and lag 1; 3-year = cumulative effect of lag 0, lag 1, and lag 2.

**North Wet**

| | SLA | | | | Flowering Time | | | |
|---|---|---|---|---|---|---|---|---|
| | **SPEI** | **CMDA** | **MAPA** | **MATA** | **SPEI** | **CMDA** | **MAPA** | **MATA** |
| **lag0** | 2.43 | 17.61 | **0.00** | 9.28 | 4.77 | 14.77 | **1.14** | 6.72 |
| **lag1** | 9.04 | 18.64 | 3.94 | 9.33 | 6.31 | 15.33 | **1.82** | 6.15 |
| **lag2** | 8.61 | 15.13 | 3.27 | 8.31 | 6.00 | 15.19 | **0.00** | 5.49 |
| **2-year** | 8.24 | 19.47 | 4.92 | 10.69 | 7.09 | 17.17 | 2.64 | 8.12 |
| **3-year** | 9.20 | 20.08 | 5.28 | 11.17 | 7.47 | 17.48 | 2.34 | 8.61 |

**Centre Wet**

| | SLA | | | | Flowering Time | | | |
|---|---|---|---|---|---|---|---|---|
| | **SPEI** | **CMDA** | **MAPA** | **MATA** | **SPEI** | **CMDA** | **MAPA** | **MATA** |
| **lag0** | 4.08 | 13.62 | **1.95** | 3.98 | 4.60 | 13.03 | **0.03** | 3.96 |
| **lag1** | 4.94 | 14.04 | **1.93** | 3.24 | 4.78 | 11.81 | **0.66** | 3.99 |
| **lag2** | 2.71 | 13.90 | **0.00** | 3.71 | 4.34 | 12.52 | **0.00** | 3.39 |
| **2-year** | 4.61 | 14.60 | 2.51 | 4.30 | 5.67 | 13.46 | **1.21** | 5.12 |
| **3-year** | 3.55 | 13.53 | **1.32** | 5.08 | 6.01 | 14.27 | **0.87** | 5.68 |

**South Wet**

| | SLA | | | | Flowering Time | | | |
|---|---|---|---|---|---|---|---|---|
| | **SPEI** | **CMDA** | **MAPA** | **MATA** | **SPEI** | **CMDA** | **MAPA** | **MATA** |
| **lag0** | 8.18 | 15.86 | 4.28 | 7.58 | 7.93 | 15.17 | 3.45 | 7.64 |
| **lag1** | 4.11 | 10.13 | **0.00** | 5.55 | 7.52 | 15.14 | 2.30 | 7.26 |
| **lag2** | 4.49 | 16.84 | **0.61** | 6.16 | 6.82 | 16.57 | **0.00** | 6.74 |
| **2-year** | 8.59 | 11.72 | 4.59 | 7.90 | 8.97 | 14.93 | 2.89 | 8.82 |
| **3-year** | 7.46 | 11.81 | 2.94 | 8.15 | 9.11 | 14.89 | **0.28** | 9.27 |

**North Dry**

| SLA | | | | Flowering Time | | | |
|---|---|---|---|---|---|---|---|
| SPEI | CMDA | MAPA | MATA | SPEI | CMDA | MAPA | MATA |
| 8.89 | 18.86 | 3.99 | 9.66 | 7.20 | 17.14 | **0.36** | 8.03 |
| 9.23 | 17.86 | 4.97 | 9.47 | 6.97 | 16.69 | **1.42** | 7.86 |
| 6.50 | 14.38 | **0.00** | 7.00 | 7.19 | 16.29 | **1.28** | 5.93 |
| 9.63 | 19.34 | 5.62 | 10.81 | 7.38 | 18.26 | **0.00** | 9.15 |
| 10.43 | 20.28 | 5.31 | 11.27 | 7.83 | 18.61 | **1.50** | 9.50 |

**Centre Dry**

| SLA | | | | Flowering Time | | | |
|---|---|---|---|---|---|---|---|
| SPEI | CMDA | MAPA | MATA | SPEI | CMDA | MAPA | MATA |
| 4.87 | 13.41 | **1.04** | 4.94 | 5.46 | 13.75 | **0.81** | 6.11 |
| 5.31 | 13.91 | **0.00** | 4.86 | 7.14 | 15.29 | 2.96 | 4.95 |
| 5.79 | 13.98 | **1.39** | 4.48 | 4.46 | 12.42 | **0.00** | 3.84 |
| 6.68 | 14.66 | 2.15 | 6.25 | 7.70 | 15.64 | 2.56 | 7.60 |
| 7.31 | 15.12 | 2.29 | 6.84 | 8.63 | 16.29 | 4.05 | 7.86 |

**South Dry**

| SLA | | | | Flowering Time | | | |
|---|---|---|---|---|---|---|---|
| SPEI | CMDA | MAPA | MATA | SPEI | CMDA | MAPA | MATA |
| 2.72 | 13.81 | **0.00** | **0.24** | 3.74 | 12.47 | **0.25** | 2.99 |
| 5.50 | 14.84 | **1.68** | 5.20 | 3.50 | 12.28 | **0.03** | 3.53 |
| 5.39 | 14.64 | **1.92** | 4.55 | 3.53 | 13.27 | **0.00** | 2.66 |
| 5.30 | 15.72 | **1.45** | 4.81 | 5.25 | 12.87 | **1.37** | 4.81 |
| 5.84 | 16.09 | 2.23 | 5.72 | 5.46 | 13.09 | **1.59** | 5.15 |

**Table S3.** Slope, p-values and $R^2$ for autocorrelation and climate average predicting trait change range-wide across the study period. SPEI = Standardised Precipitation Evapotranspiration Index; CMDA = Hargreaves Climate Moisture Deficit Anomaly; MAPA = log Mean Annual Precipitation Anomaly; MATA = Mean Annual Temperature Anomaly. No p-values are significant after correcting for multiple tests.

| | Autocorrelation | | | Mean Climate Change | | |
|---|---|---|---|---|---|---|
| | **slope** | **p-value** | **$R^2$** | **slope** | **p-value** | **$R^2$** |
| **SLA - Wet** | | | | | | |
| **CMDA** | -37.22 | 0.46 | 0.08 | 0.21 | 0.47 | 0.08 |
| **MAPA** | 36.76 | 0.47 | 0.16 | -69.94 | 0.63 | 0.16 |
| **MATA** | -8.43 | 0.89 | 0.00 | 1.80 | 0.97 | 0.00 |
| **SLA - Dry** | | | | | | |
| **CMDA** | 22.97 | 0.70 | 0.06 | -0.26 | 0.45 | 0.06 |
| **MAPA** | 103.15 | 0.086 | 0.30 | 117.59 | 0.47 | 0.30 |
| **MATA** | 30.12 | 0.67 | 0.06 | -43.29 | 0.48 | 0.06 |
| **Date of Flowering - Wet** | | | | | | |
| **CMDA** | -10.33 | 0.20 | 0.20 | 0.00 | 0.95 | 0.20 |
| **MAPA** | 1.05 | 0.89 | 0.32 | -35.30 | 0.13 | 0.32 |
| **MATA** | -9.17 | 0.32 | 0.23 | -2.95 | 0.70 | 0.23 |
| **Date of Flowering - Dry** | | | | | | |
| **CMDA** | 1.91 | 0.80 | 0.03 | 0.01 | 0.78 | 0.03 |
| **MAPA** | -8.80 | 0.27 | 0.14 | -18.79 | 0.41 | 0.14 |
| **MATA** | -11.63 | 0.17 | 0.24 | 10.22 | 0.16 | 0.24 |

**Table S4**. Summary statistics for auto-correlation and climate average predicting trait change range-wide for 5-time chunks during the timeseries. CMDA = Hargreaves Climate Moisture Deficit Anomaly; MAPA = log Mean Annual Precipitation Anomaly; MATA = Mean Annual Temperature Anomaly. No p-values are significant after correcting for multiple tests.

| | Autocorrelation | | | Mean Climate Change | | |
|---|---|---|---|---|---|---|
| | **slope** | **p-value** | **$R^2$** | **slope** | **p-value** | **$R^2$** |
| **SLA - Wet** | | | | | | |
| **CMDA** | -15.82 | 0.44 | 0.04 | 0.04 | 0.54 | 0.04 |
| **MAPA** | 4.08 | 0.82 | 0.02 | -34.38 | 0.48 | 0.02 |
| **MATA** | 10.20 | 0.66 | 0.03 | 7.07 | 0.39 | 0.03 |
| **SLA - Dry** | | | | | | |
| **CMDA** | -20.32 | 0.54 | 0.02 | 0.04 | 0.74 | 0.02 |
| **MAPA** | 9.23 | 0.75 | 0.00 | -5.25 | 0.95 | 0.00 |
| **MATA** | -3.66 | 0.92 | 0.02 | 5.76 | 0.67 | 0.02 |
| **Date of Flowering - Wet** | | | | | | |
| **CMDA** | 0.37 | 0.88 | 0.03 | -0.01 | 0.39 | 0.03 |
| **MAPA** | 0.83 | 0.68 | 0.05 | -6.11 | 0.28 | 0.05 |
| **MATA** | 1.20 | 0.66 | 0.03 | -0.29 | 0.76 | 0.03 |
| **Date of Flowering - Dry** | | | | | | |
| **CMDA** | -0.94 | 0.63 | 0.02 | 0.00 | 0.54 | 0.02 |
| **MAPA** | 1.26 | 0.46 | 0.03 | 2.13 | 0.64 | 0.03 |
| **MATA** | -4.66 | 0.02 | 0.25 | -0.16 | 0.82 | 0.25 |

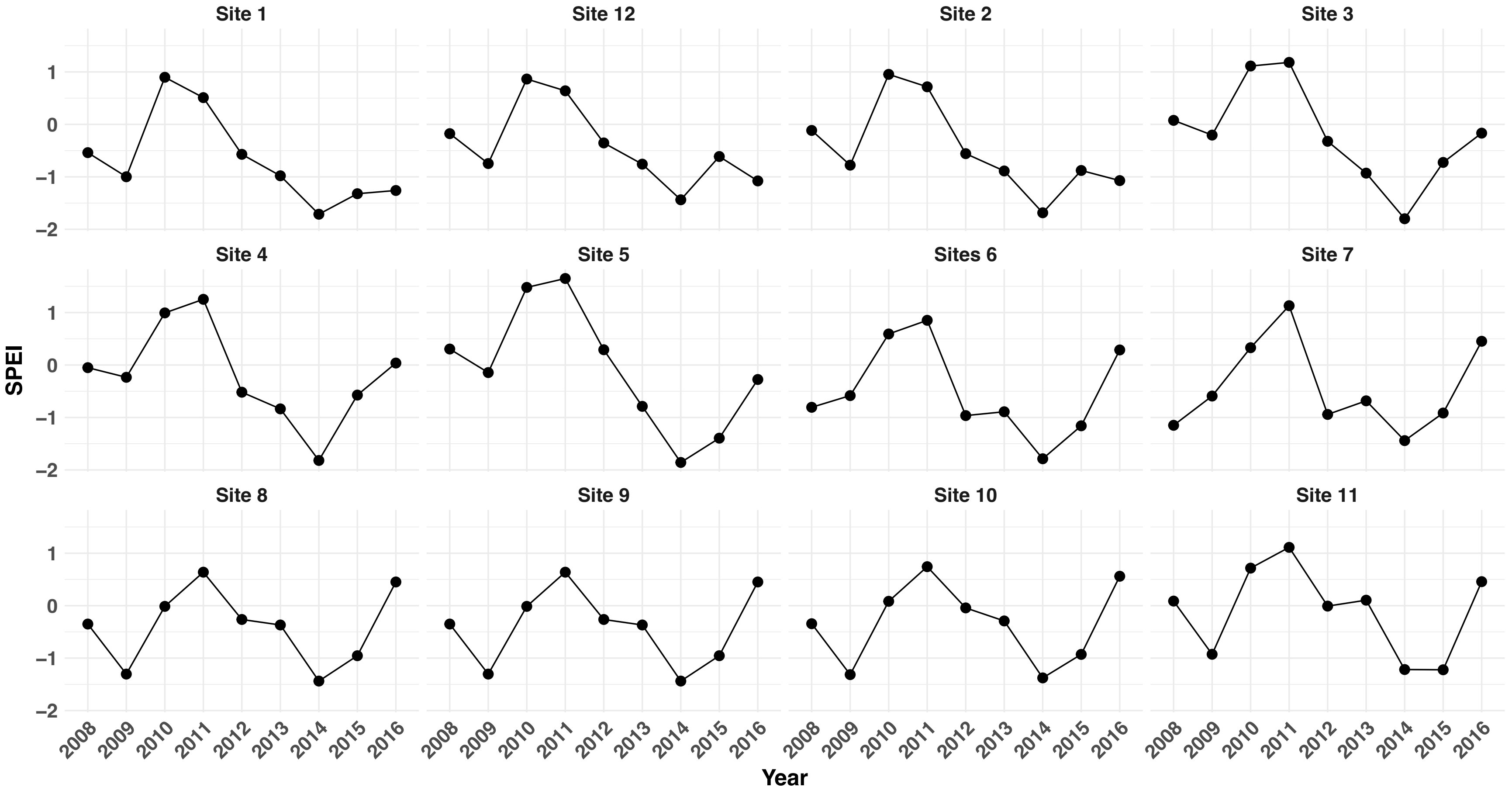


**Figure S1.** SPEI (Standardized Precipitation Evapotranspiration Index) values for water years 2008-2016 across 12 *M. cardinalis* site in California and Oregon.

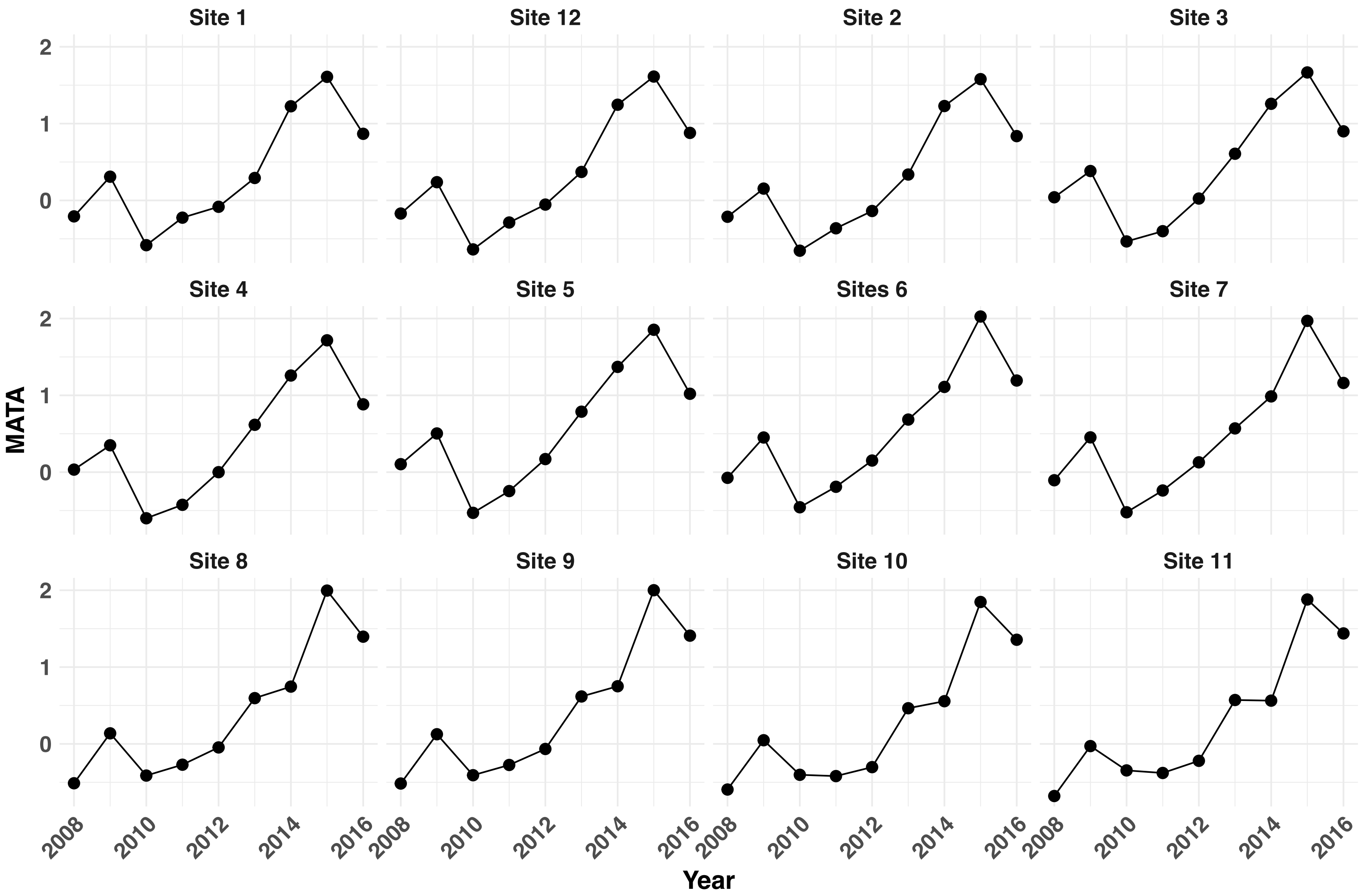


**Figure S2**. MATA (mean annual temperature anomaly) values for water years 2008-2016 across 12 *M. cardinalis* site in California and Oregon.

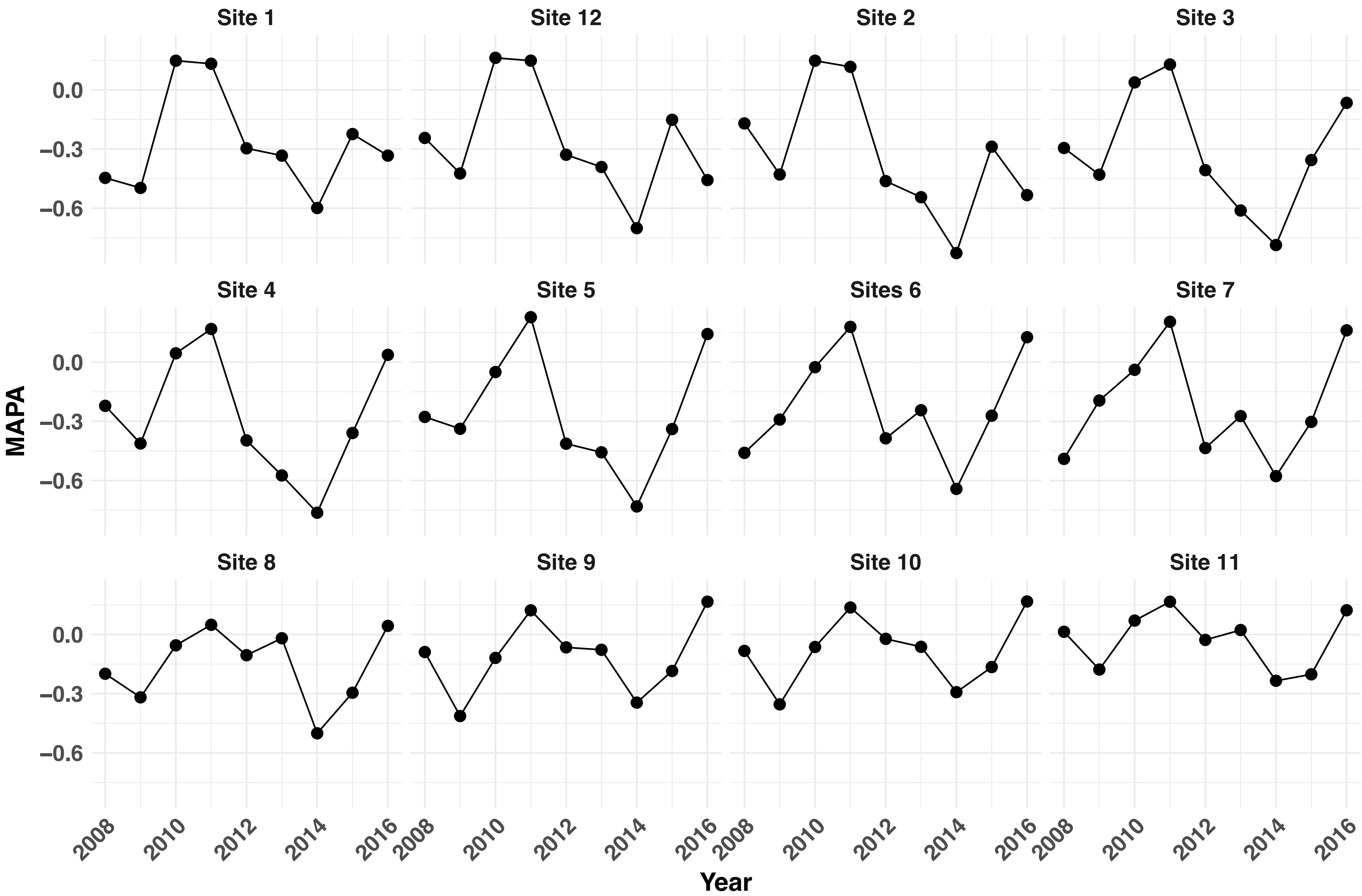


**Figure S3**. MAPA (mean annual precipitation anomaly) values for water years 2008-2016 across 12 *M. cardinalis* site in California and Oregon.

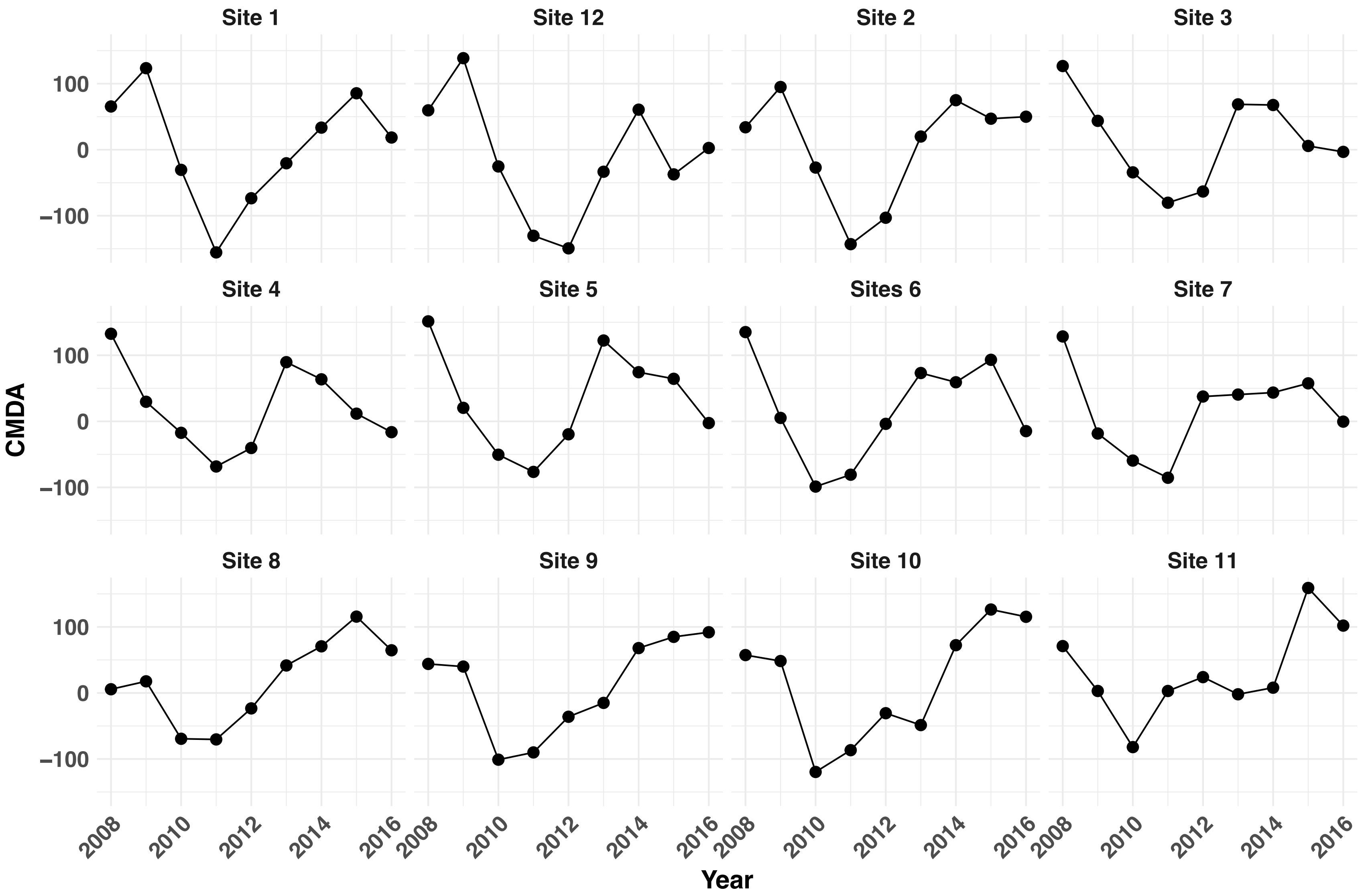


**Figure S4**. CMDA (cumulative moisture deficit anomaly) values for water years 2008-2016 across 12 *M. cardinalis* site in California and Oregon.

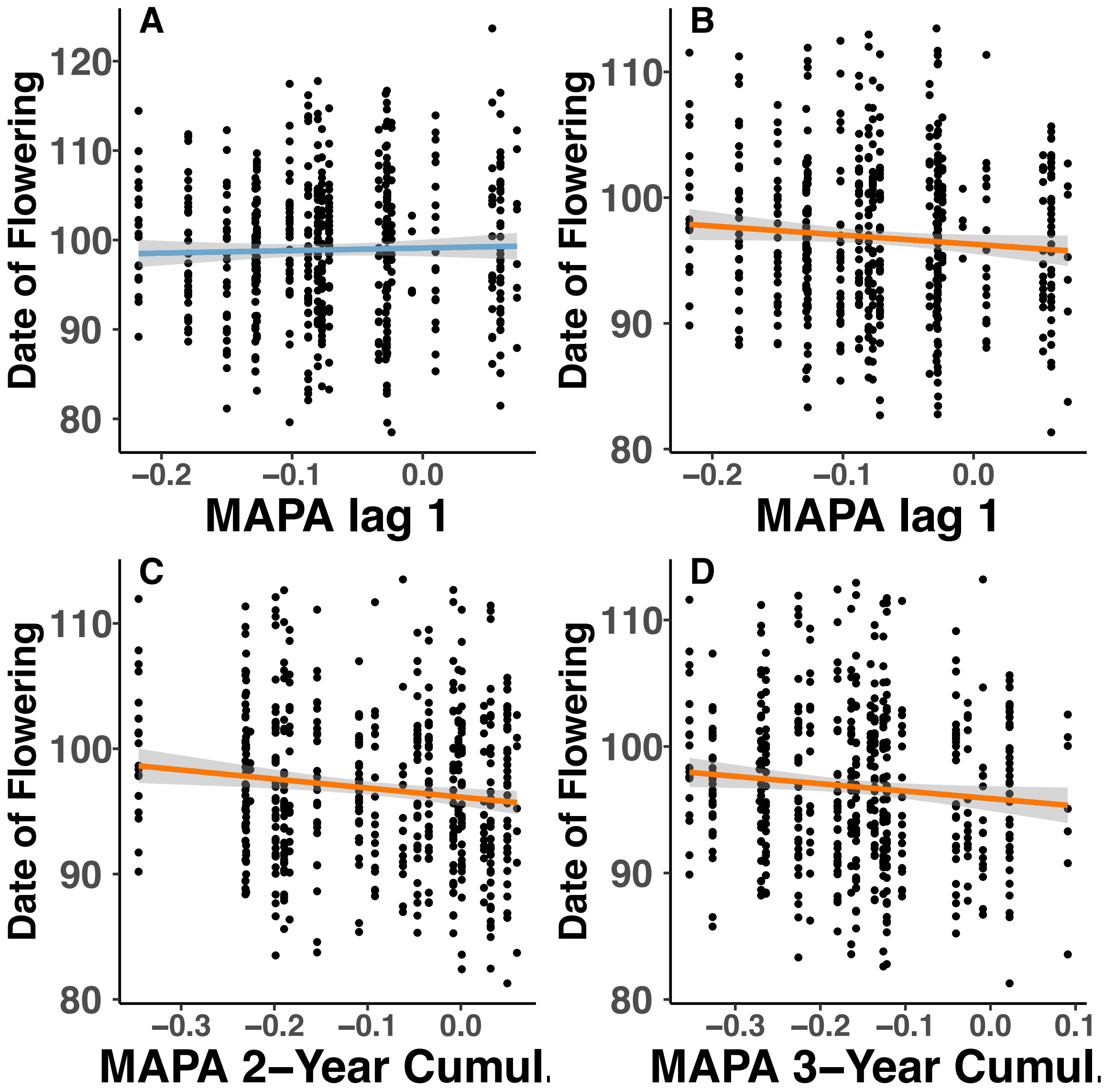


**Figure S5. Evolution of Specific Leaf Area (SLA) and date of flowering across mean annual precipitation anomaly (MAPA) in the northern part of the *Mimulus cardinalis* range.** Date of flowering is explained by A) lag 1 when grown under well-watered conditions. Date of flowering is also explained by B) lag 1, C) 2-year cumulative MAPA and D) 3-year cumulative MAPA when grown under drought conditions. Lag 1 = effect of climate from one year prior. Blue regression line = well-watered treatment; orange regression line = drought treatment.

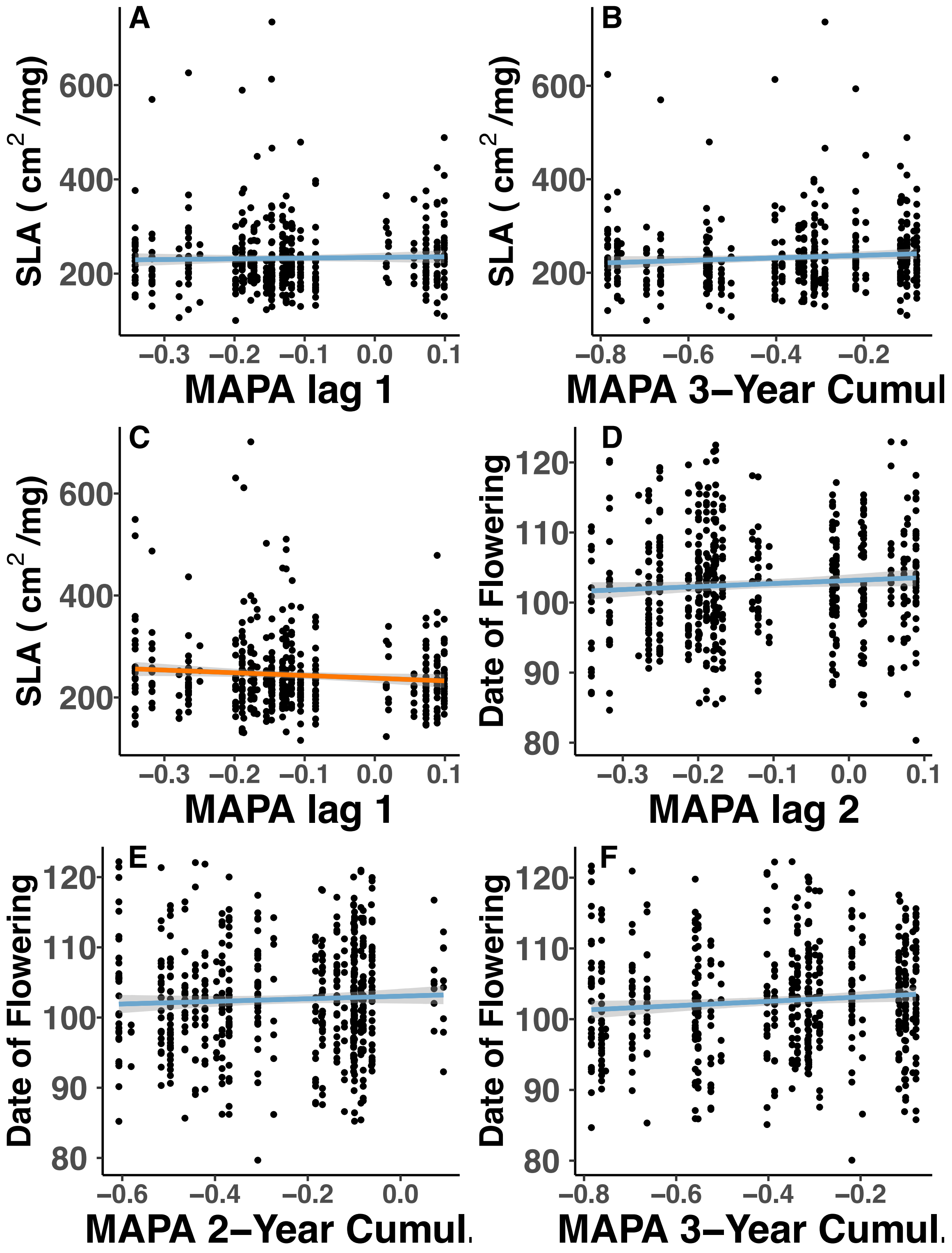


**Figure S6. Evolution of Specific Leaf Area (SLA) and date of flowering across mean annual precipitation anomaly (MAPA) in the central part of the *Mimulus cardinalis* range.** A) SLA is explained by lag 0 and B) 3-year cumulative MAPA when grown under a well-watered treatment. C) SLA is also explained by MAPA lag 1 when plants grown under a drought treatment. Date of flowering is explained by D) lag 2, E) 2-year cumulative MAPA, F) 3-year cumulative MAPA when grown under a well-watered treatment. Lag 0 = effect of current year's climate; lag 1 = effect of climate from one year prior; lag 2 = effect of climate from two years prior. Blue regression line = well-watered treatment; orange regression line = drought treatment.

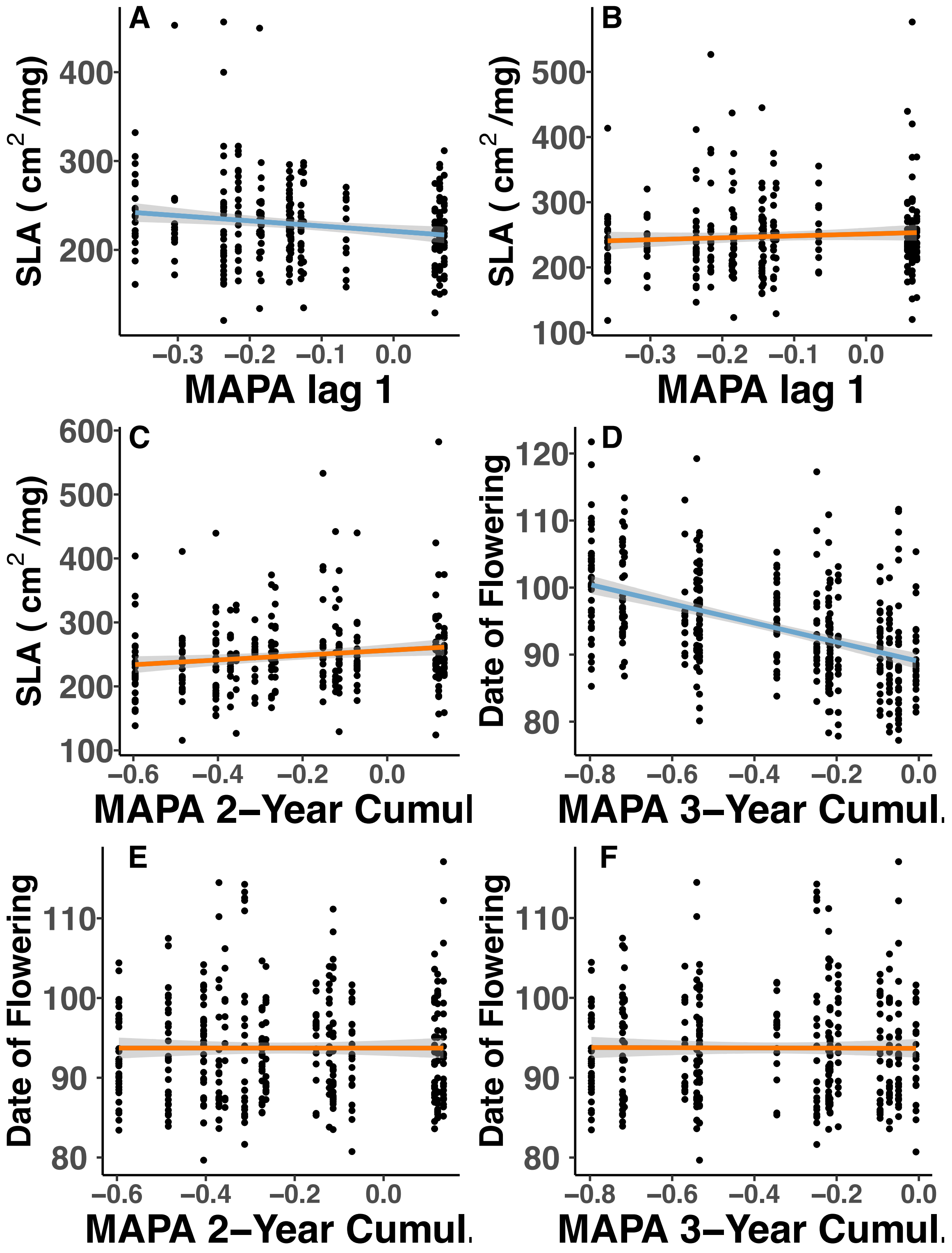


**Figure S7. Evolution of Specific Leaf Area (SLA) and date of flowering across mean annual precipitation anomaly (MAPA) in the southern part of the *Mimulus cardinalis* range.** A)

SLA is explained by lag 1 when grown under well-watered conditions. SLA is also explained by B) lag 1, C) 2-year cumulative MAPA, and D) 3-year cumulative MAPA when grown under a drought treatment. Flowering time is explained by E) 2-year cumulative MAPA, and F) 3-year cumulative MAPA when grown under a drought treatment. Lag 1 = effect of climate from one year prior; lag 2 = effect of climate from two years prior. Blue regression line = well-watered treatment; orange regression line = drought treatment.

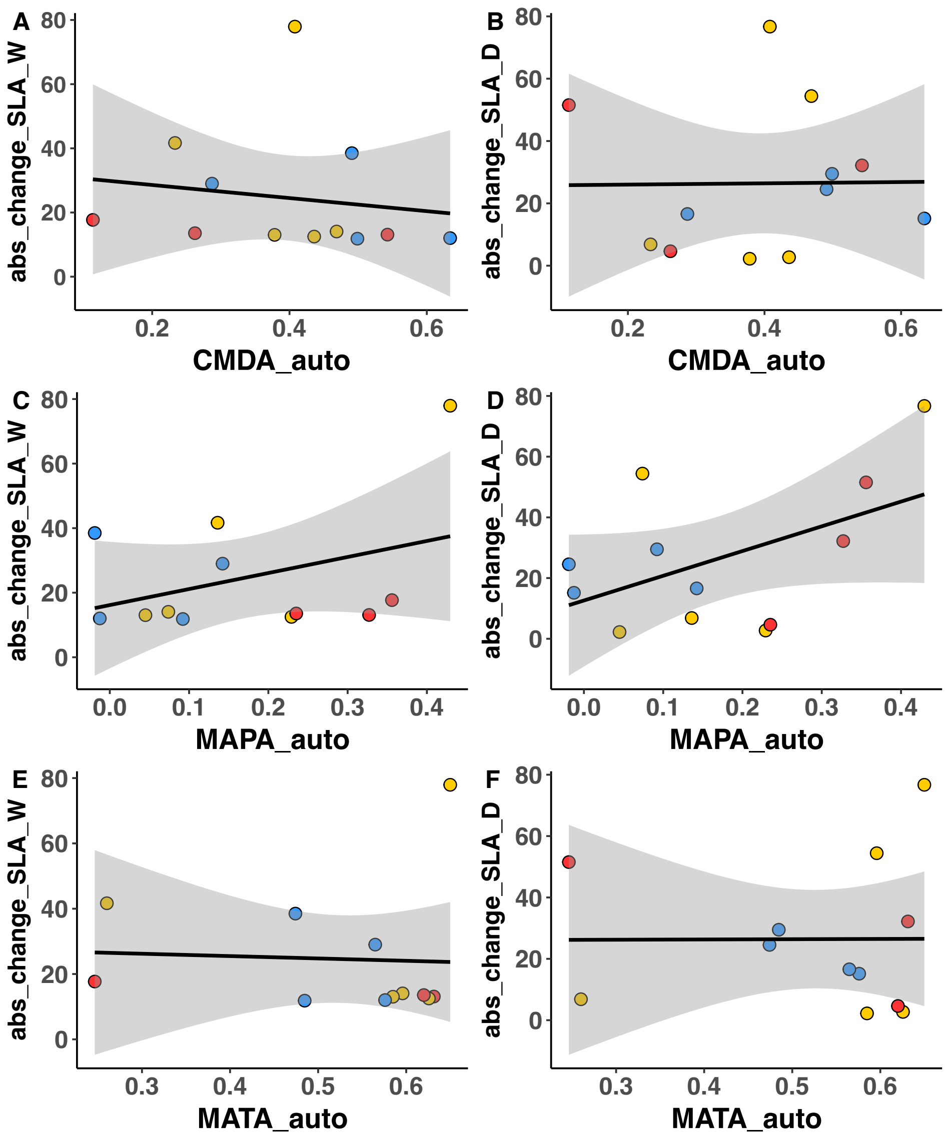


**Figure S8** Environmental autocorrelation from 2010 to 2016 predicting the absolute change in SLA under (A,C,E) wet and (B,D,F) dry experimental treatments. Red point = population from Southern Region; yellow points population from Central Region; blue point population from Northern Region.

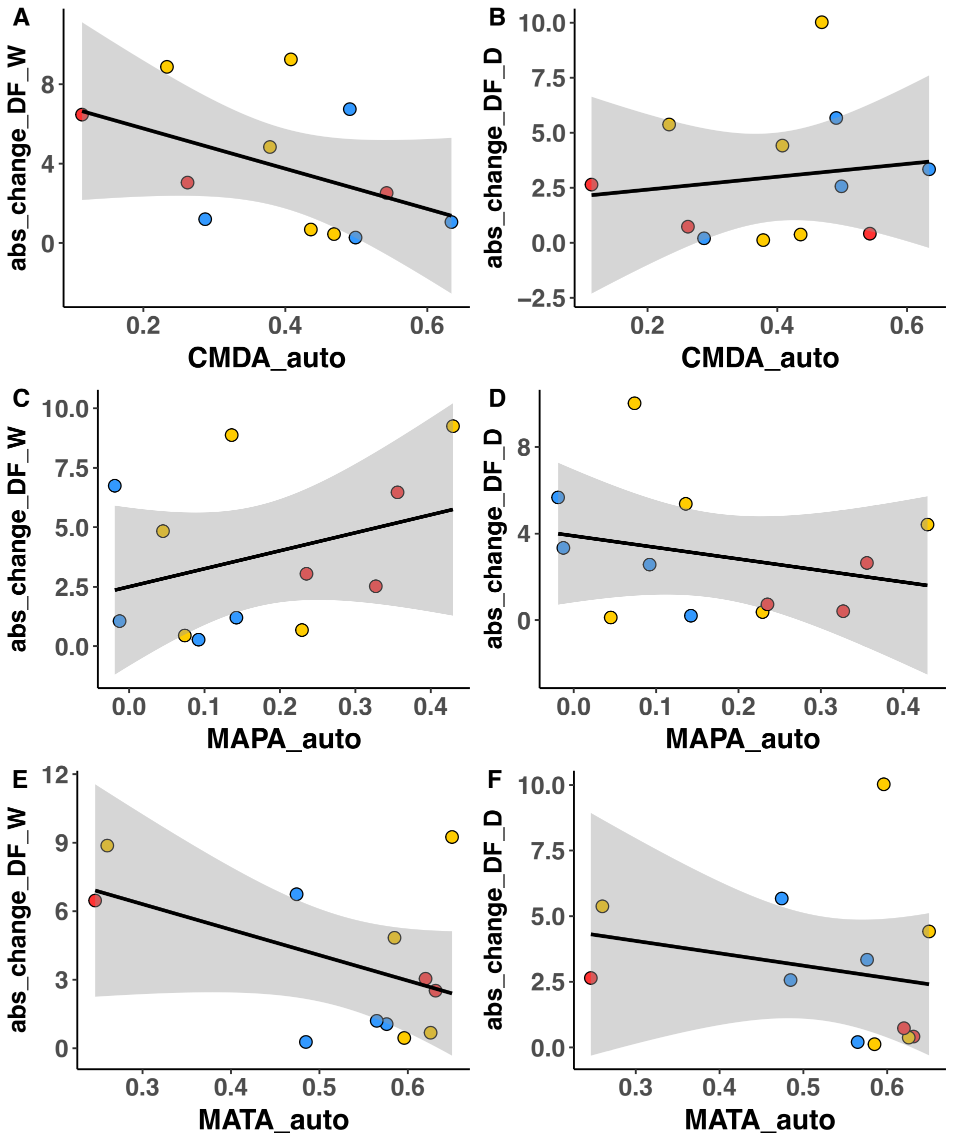


**Figure S9** Environmental autocorrelation from 2010 to 2016 predicting the absolute change in flowering date under (A,C,E) wet and (B,D,F) dry experimental treatments. Red point = population from Southern Region; yellow points population from Central Region; blue point population from Northern Region.

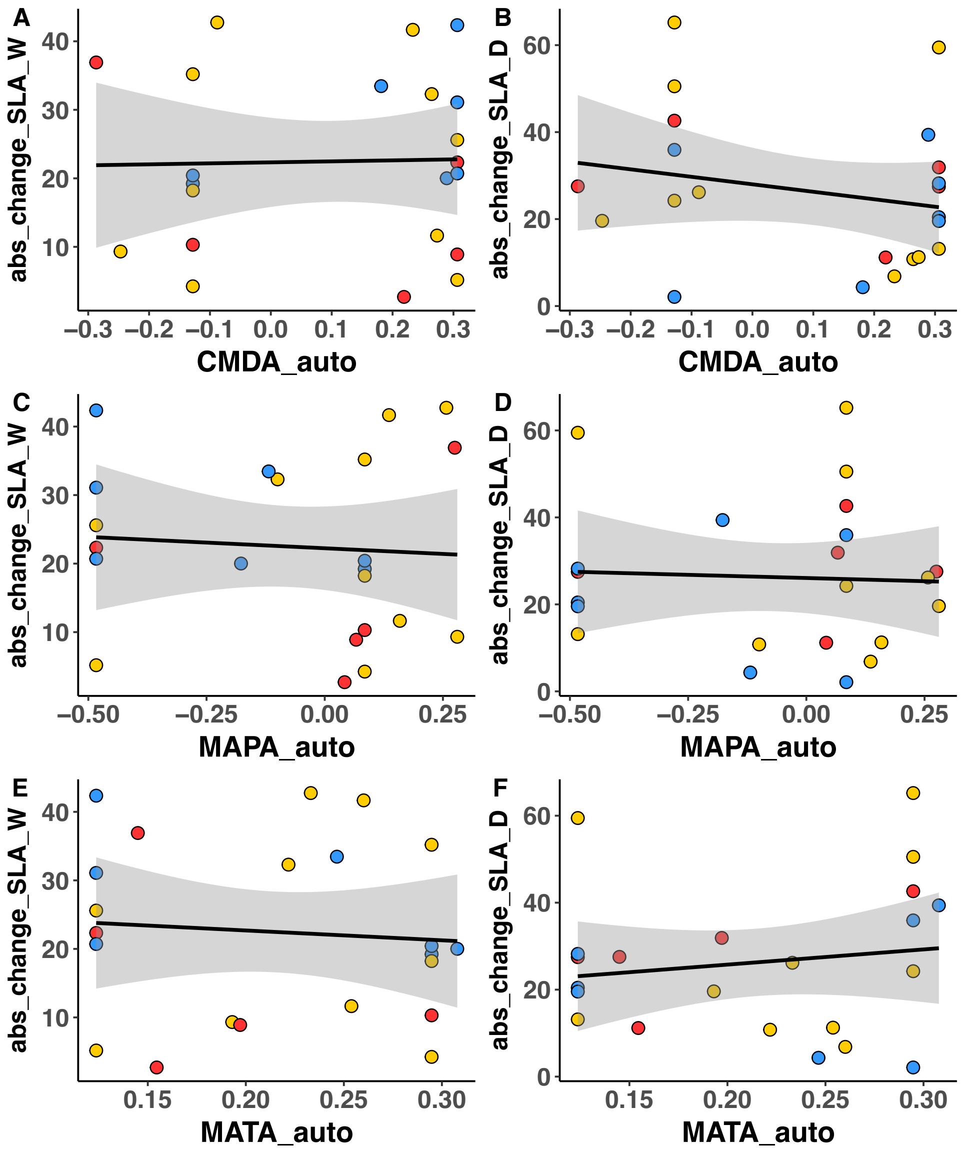


**Figure S10** Environmental autocorrelation under five 3-year periods from 2010 to 2016 predicting the absolute change in SLA under (A,C,E) wet and (B,D,F) dry experimental treatments. Red point = population from Southern Region; yellow points population from central region; blue point population from Northern Region.

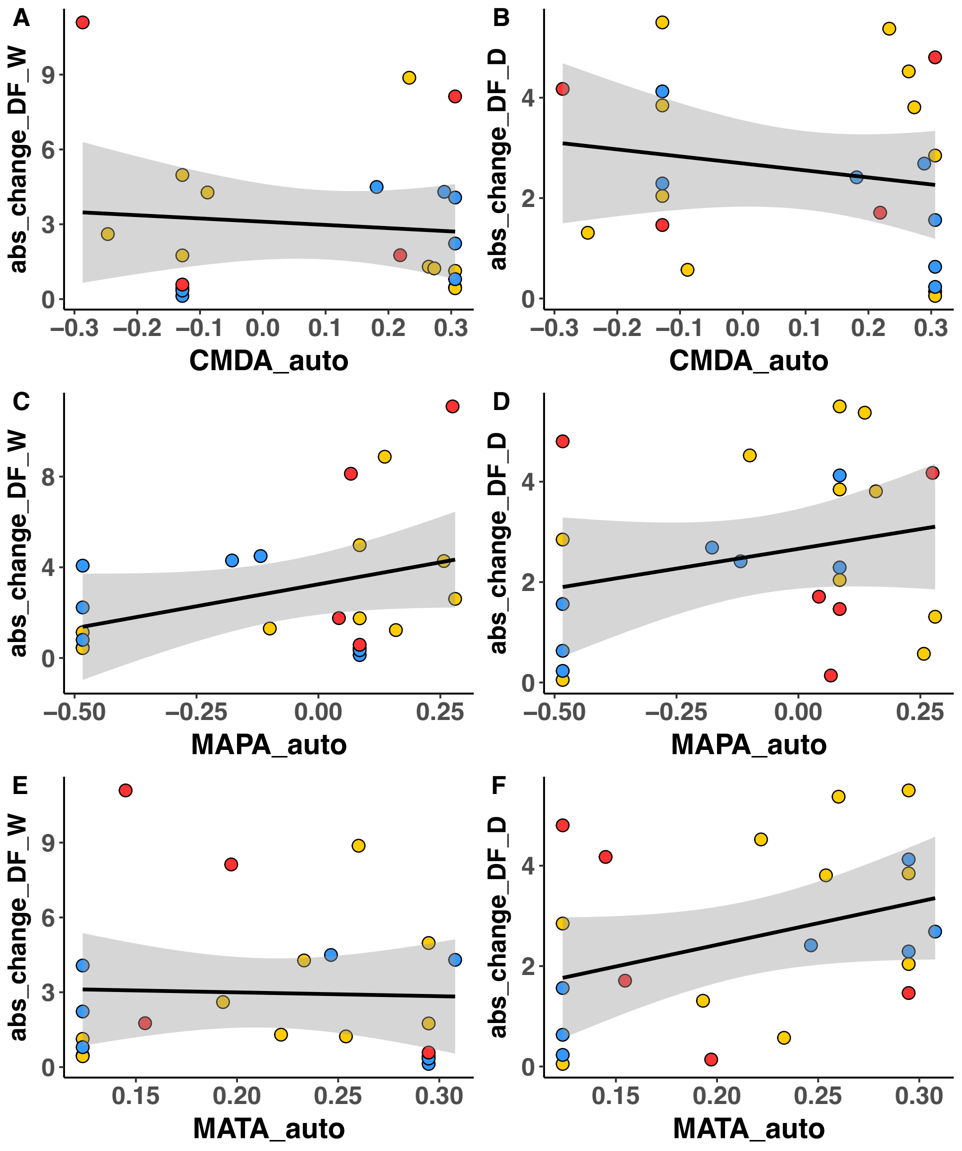


**Figure S11** Environmental autocorrelation under five 3-year periods from 2010 to 2016 predicting the absolute change in date of lowering under (A,C,E) wet and (B,D,F) dry experimental treatments. Red point = population from Southern Region; yellow points population from Central Region; blue point population from Northern Region.
